# The lymphangiogenic Ccbe1-Adamts-Vegfc axis shapes cerebrovascular regionalization via selective patterning of fenestrated vascular beds

**DOI:** 10.64898/2026.10.14.759138

**Authors:** Luke D. Buck, Vani Thakur, Alyssa G. Storm, Fatma Z. Bozdag, Nathanael J. Lee, Jun Xiong Leong, Ella N. Kazazic, Amanda E. Lam, Angelene A. Getz, Ryota L. Matsuoka

**Affiliations:** Department of Neurosciences, Cleveland Clinic Research, Cleveland, USA; Department of Molecular Medicine, Cleveland Clinic Lerner College of Medicine, Case Western Reserve University, Cleveland, USA

**Author notes:** Corresponding author: Ryota L. Matsuoka.

**Keywords:** brain angiogenesis, cerebrovascular heterogeneity, lymphangiogenic signaling, fenestrated vessels, blood-brain barrier, zebrafish

## Abstract

Organ development and function require regional specialization of vascular networks, yet the underlying molecular mechanisms remain poorly understood. Here, we identify a non-canonical role for the lymphangiogenic Ccbe1-Adamts-Vegfc axis in shaping cerebrovascular regionalization in zebrafish. A *ccbe1* knock-in reporter revealed highly restricted expression in the developing brain, with its genetic loss causing early vascularization defects at the pineal gland-choroid plexus interface. Simultaneous Vegfa signaling deficiency markedly exacerbated these regional vascular defects, accompanied by disrupted blood flow, local hemorrhage, and choroid plexus epithelial disorganization, while largely preserving neighboring barrier-forming vessels. Loss of *vegfc* closely phenocopied *ccbe1* mutant defects, with fenestrated vascular patterning requiring both Vegfr2 and Vegfr3 signaling, whereas disruption of *adamts3* and *adamts14* indicated region-specific requirements for Vegfc-activating proteases. Conversely, pineal gland-specific overexpression of activated Vegfc partially rescued the vascularization defects. Collectively, these findings suggest that spatially restricted Ccbe1-Adamts-Vegfc activity provides angiogenic signaling specificity enabling fenestrated cerebrovascular patterning and regionalization.

**Highlights:**

- The lymphangiogenic Ccbe1-Adamts-Vegfc axis patterns fenestrated brain vasculature
- Vegfc and Vegfa signaling cooperate to promote regional cerebrovascular development
- Combined Vegfc and Vegfa pathway disruptions alter local cerebrovascular function
- Localized lymphangiogenic signaling rescues fenestrated brain vascular defects

## Introduction

The unique functions of individual organs depend on organotypic vascular networks established during embryogenesis. How diverse vascular endothelial cell (EC) identities and properties are generated within and across organs remains an important question in vascular biology and organ physiology. The brain is distinguished from peripheral organs by highly specialized vascular barrier properties, collectively known as the blood-brain barrier (BBB) (*1, 2*). However, diverse brain functions also require region-specific cerebrovascular beds that lack BBB properties, creating considerable intra-organ vascular heterogeneity (*3, 4*). Despite substantial progress in understanding the cellular and molecular mechanisms of BBB specialization and integrity, how fenestrated permeable capillaries in the choroid plexuses (CPs) and circumventricular organs (CVOs) form remains poorly understood. Given the essential roles of both barrier-forming and fenestrated vascular beds in brain function and homeostasis, it is crucial to define their developmental mechanisms underlying cerebrovascular regionalization.

Fenestrated capillaries in the CPs and CVOs support brain development, physiology, and immunity by facilitating molecular exchange between neural tissues and the circulation, while also contributing to fluid homeostasis and immune cell trafficking (*5, 6*). Moreover, the CP has received increasing attention in the context of neurological diseases, due to its vital roles in forming the blood-cerebrospinal fluid (CSF) barrier and regulating CSF production and clearance (*7–10*). Likewise, as a neuroendocrine CVO, the pineal gland relies on fenestrated vasculature for efficient communication with the circulation and for regulation of circadian rhythms through melatonin secretion (*3, 11*). However, despite their critical physiological roles, the formation of these fenestrated cerebrovascular beds remains largely unknown.

We recently demonstrated that vascularization of both the zebrafish CP and PG requires coordinated angiogenic (Vegfa) and lymphangiogenic (Vegfc/d) signaling (*12–14*). Specifically, multiple Vegf ligands and their receptors participate in regulating these developmental processes, indicating functional interplay between angiogenic and lymphangiogenic pathways. Because blood and lymphatic ECs typically exhibit distinct molecular identities and developmental programs (*15–18*), the requirement for lymphangiogenic signaling during cerebrovascular development was unexpected. These findings raised a fundamental question: how does a lymphangiogenic signaling module operate in patterning fenestrated cerebrovascular beds without affecting neighboring BBB-type vessels?

Unlike VEGF-A, whose signaling properties are regulated in part through alternative splicing (*19*), VEGF-C is produced as an inactive propeptide that requires extracellular proteolytic processing for activation (*20*). Proteolytic processing enables high-affinity binding of VEGF-C to VEGFR3 and VEGFR2, thereby stimulating downstream signaling cascades (*20, 21*). The secreted protein CCBE1 acts as a critical cofactor that facilitates proteolytic activation of VEGF-C by ADAMTS metalloproteinases (*22–26*). Accordingly, genetic loss of *Vegfc* or *Ccbe1* results in nearly identical lymphangiogenic defects across species (*22, 24, 27, 28*), demonstrating the evolutionary conservation of this signaling axis.

Given the continuous, interconnected nature of the cerebrovasculature, how bioactive signaling events are spatiotemporally coordinated to generate cerebrovascular regional heterogeneity remains an important question. Despite reported roles for the Ccbe1-Adamts-Vegfc axis in lymphatic and coronary vessel development (*29–31*), its contribution to cerebrovascular formation remains largely unexplored. Since VEGF-C activation requires extracellular proteolytic machinery, its activating components could contribute to local restriction of VEGF-C activity during brain vascularization. We therefore hypothesized that spatiotemporal regulation of VEGF-C activation allows the signaling specificity required for patterning fenestrated cerebrovascular beds. Here, using newly developed zebrafish knock-in reporters, genetic analyses, and cell type-specific approaches, we show spatial and functional requirements of the Ccbe1-Adamts-Vegfc axis during fenestrated cerebrovascular development.

## Results

### A novel knock-in zebrafish reporter reveals prominent, localized *ccbe1* expression in brain regions associated with fenestrated vasculature

To characterize endogenous *ccbe1* expression patterns in live zebrafish and investigate its potential roles during brain vascularization, we generated a new knock-in line in which the zebrafish codon-optimized Gal4 derivative KalTA4 was integrated near the stop codon of the endogenous *ccbe1* coding sequence using CRISPR/Cas9-mediated genome targeting (Fig. 1A). An E2A bicistronic peptide sequence was incorporated to enable expression of KalTA4 from the endogenous *ccbe1* transcript. Both heterozygous and homozygous knock-in animals carrying UAS-driven fluorescent reporters exhibited robust, spatially restricted reporter expression (Fig. 1B, C). Previous BAC transgenic and knock-in reporter studies have characterized *ccbe1* expression in the developing trunk (*32, 33*). Consistent with these studies, our knock-in reporter reproduced previously described trunk expression patterns (Fig. 1B, C), supporting its reliability for visualizing endogenous *ccbe1* expression.

**Fig. 1.**
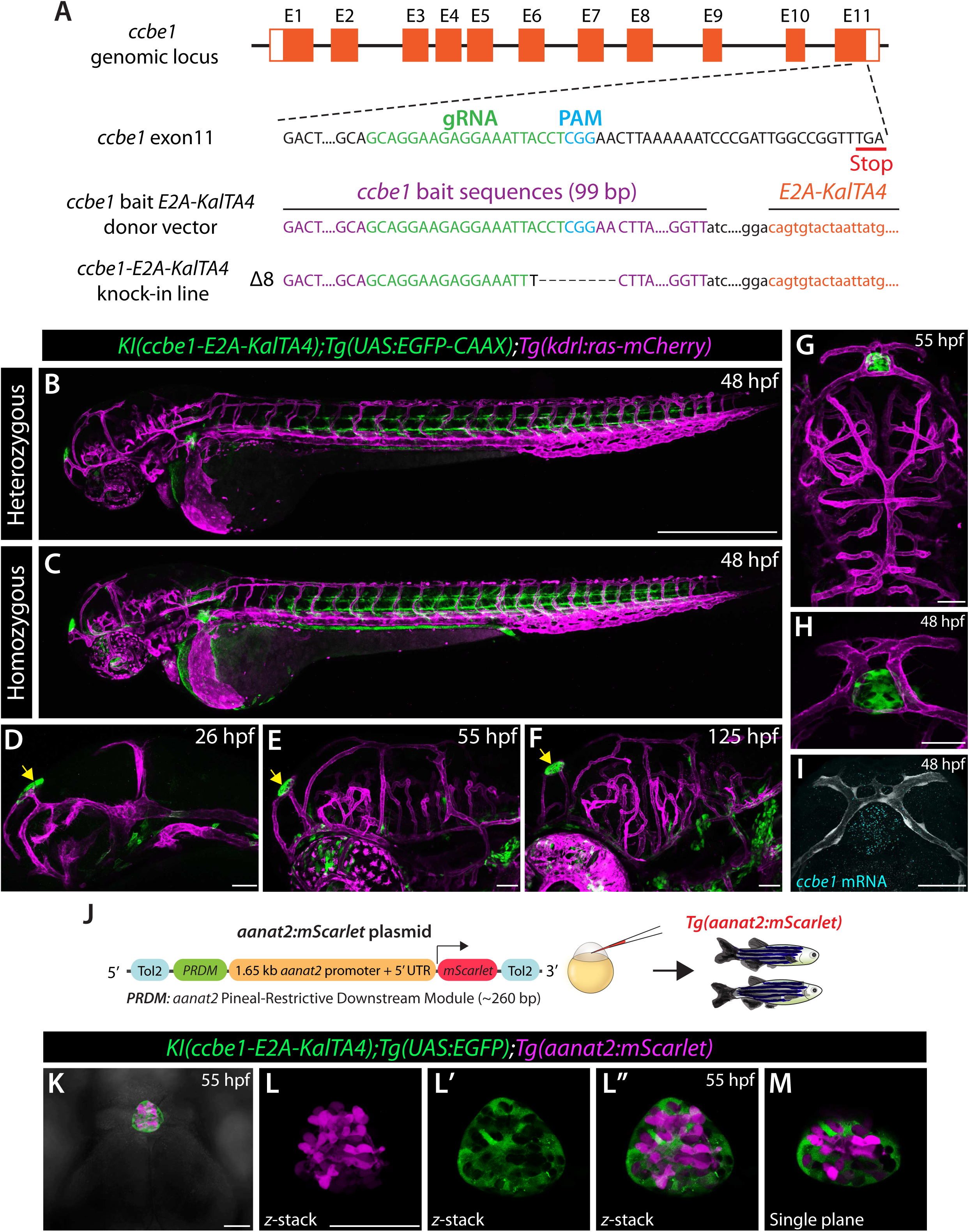
Generation of a novel knock-in zebrafish reporter reveals prominent, localized *ccbe1* expression in the developing pineal gland. (**A**) Strategy for generating a *ccbe1* knock-in zebrafish line and genomic characterization of the established line. (**B, C**) Lateral view of 48 hpf *ccbe1* knock-in heterozygous and homozygous whole embryos carrying the *Tg(UAS:EGFP-CAAX);Tg(kdrl:ras-mCherry)* reporters, showing similar EGFP expression domains with the homozygous embryo exhibiting brighter expression. (**D**-**F**) Developmental characterization of 26, 55, and 125 hpf heterozygous *ccbe1* knock-in fish carrying the *Tg(UAS:EGFP-CAAX);Tg(kdrl:ras-mCherry)* reporters, showing prominent EGFP expression in the pineal gland (yellow arrows). (**G**) Dorsal view of 55 hpf *ccbe1* knock-in heterozygous embryo carrying the *Tg(UAS:EGFP-CAAX);Tg(kdrl:ras-mCherry)* reporters, showing strong EGFP signals localized to the pineal gland. (**H, I**) Side-by-side comparison of a 48 hpf heterozygous *ccbe1* knock-in embryo carrying the *Tg(UAS:EGFP-CAAX);Tg(kdrl:ras-mCherry)* reporters (**H**) and *ccbe1* mRNA expression (**I**), showing consistent positive signals in the pineal gland. (J) Schematic diagram of the donor plasmid used to generate the *Tg(aanat2:mScarlet)* zebrafish line. (K) Dorsal view of a *Tg(aanat2:mScarlet)* embryo carrying the heterozygous *ccbe1* knock-in allele and *Tg(UAS:EGFP)*, showing specific pineal gland expression from both lines. (**L**-**M**) Magnified maximal *z*-stack projection images of a *Tg(aanat2:mScarlet)* embryo carrying the heterozygous *ccbe1* knock-in allele and *Tg(UAS:EGFP)* (**L**-**L”**) and a corresponding single-plane image (**M**), showing non-overlapping expression domains of the two reporters. Scale bars: 500 µm in **B** for **B, C**; 50 µm in **D**-**I, K**; **L** for **L’**-**L”, M**.

We next examined *ccbe1* expression during brain vascular development, where its cellular distribution and function remain poorly characterized. Strikingly, *ccbe1* reporter expression in the developing brain exhibited highly localized patterns, with the most prominent expression observed in the pineal gland (Fig. 1D-H). Reporter-positive cells were already evident in the pineal gland by 26 hours post fertilization (hpf), and this prominent expression persisted through later developmental stages. To validate these expression patterns, we examined endogenous *ccbe1* transcripts using RNAscope *in situ* hybridization. Prominent endogenous *ccbe1* mRNA signals were indeed detected in the pineal gland and showed a spatial distribution consistent with the knock-in reporter expression (Fig. 1H, I), confirming that the reporter reliably reflects endogenous *ccbe1* localization.

The prominent and persistent *ccbe1* expression within the pineal gland motivated us to investigate the cellular identity of this population. The pineal gland of non-mammalian vertebrates is recognized as a photoreceptive neuroendocrine organ that contains an intrinsic circadian oscillator and regulates daily and seasonal rhythms (*34*). Major cell populations characterized in the teleost pineal gland include photoreceptors, projection neurons, and interstitial cells with glial characteristics (*35–37*). To determine whether *ccbe1*-expressing cells represent pineal photoreceptors, we generated a transgenic reporter in which mScarlet expression is driven by regulatory elements of *aanat2* (Fig. 1J), a gene prominently expressed in pineal photoreceptors and involved in melatonin synthesis (*36*). The resulting transgenic line exhibited strong mScarlet expression specifically within the pineal gland (Fig. 1K). Simultaneous visualization of the *ccbe1* knock-in and *aanat2* transgenic reporters revealed that the two cell populations were spatially intermingled throughout the pineal gland with little cellular overlap (Fig. 1L-L’’, M). Thus, *ccbe1* expression marks a distinct cellular population within the developing pineal gland that is largely separate from *aanat2*-positive photoreceptors.

### Genetic loss of *ccbe1* causes a substantial delay in early vascularization at the pineal gland-CP interface associated with fenestrated vessels

The prominent and localized expression of *ccbe1* in the pineal gland raised the possibility that Ccbe1 contributes to vascular development in this region. The zebrafish brain contains two anatomically distinct CPs, the diencephalic and myelencephalic CP (dCP and mCP), corresponding to the third and fourth ventricular CPs in mammals, respectively (*38*). Notably, the pineal gland and dCP develop in close anatomical proximity, forming a tissue interface associated with specialized fenestrated vasculature. We therefore examined vascular development at this interface and determined how it is affected by loss of *ccbe1*.

Vascularization of the pineal gland-dCP interface begins during early embryonic development. At approximately 20 hpf, vascular sprouts emerged from the bilateral primordial midbrain channels and extended toward the pineal gland and developing dCP (Fig. 2A). These sprouts turned rostrally along the pineal gland and reached the vicinity of the developing dCP by approximately 21.5 hpf (Fig. 2B). Between 21.5 and 23 hpf, the vessels extended further ventrolaterally to establish connections with the cranial division of the primitive internal carotid artery (Fig. 2C), rapidly forming a vascular network at the pineal gland-dCP interface (Fig. 2D).

**Fig. 2.**
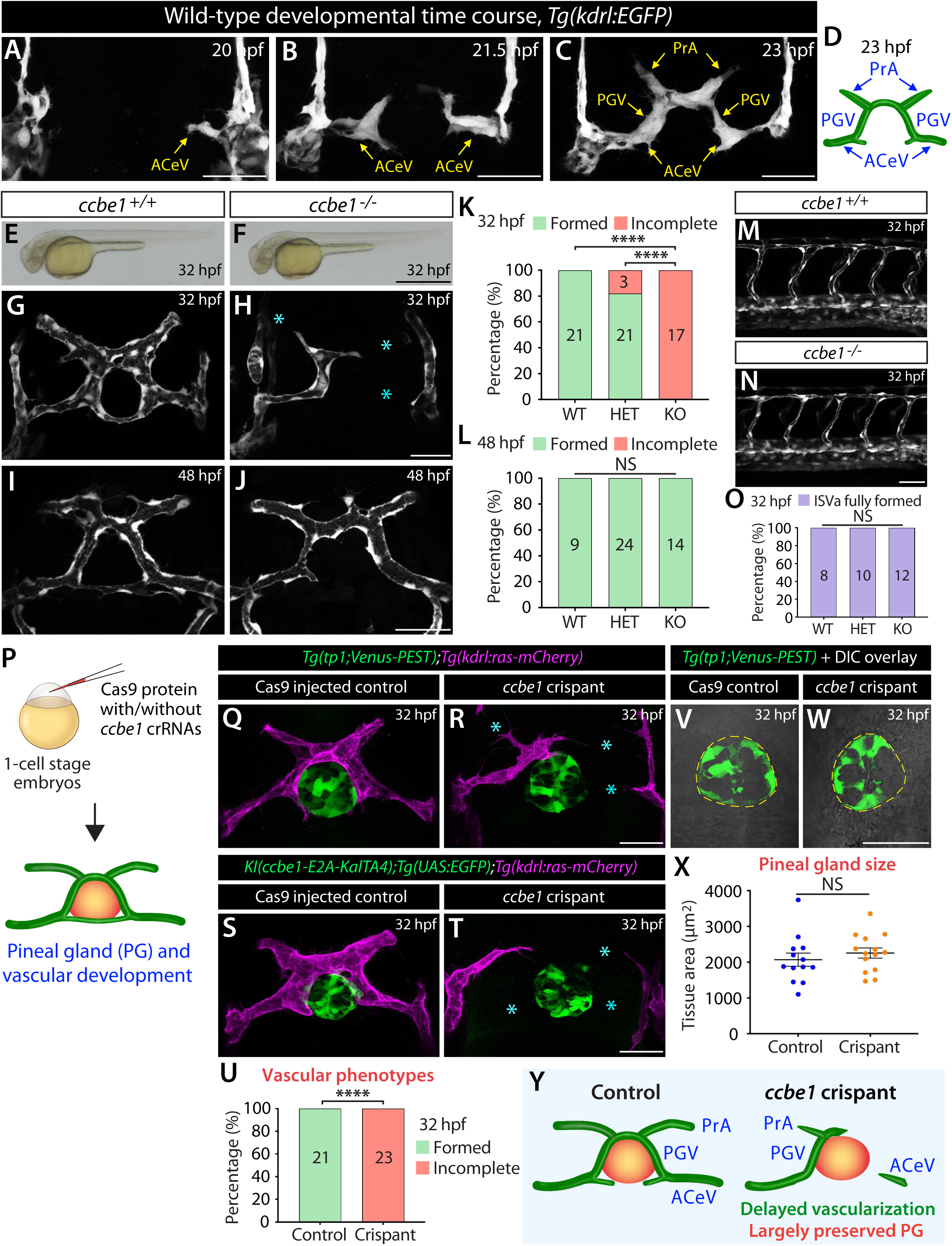
Genetic loss of *ccbe1* causes a substantial delay in the vascularization of pineal gland-diencephalic choroid plexus (dCP) interface while largely preserving pineal gland development. (A-C) Dorsal views of 20 (A), 21.5 (B), and 23 (C) hpf *Tg(kdrl:EGFP)* embryos show the rapid progression of vascularization at the pineal gland-dCP interface. PrA: prosencephalic artery, PGV: pineal gland vessel, ACeV: anterior cerebral vein. (D) Schematic diagram illustrating the wild-type (WT) vascular anatomy at the pineal gland-dCP interface at 23 hpf. (**E, F**) Lateral brightfield views of 32 hpf *ccbe1^+/+^* and *ccbe1^-/-^* embryos, showing comparable overall embryonic morphology. (**G, H**) Dorsal views of 32 hpf *ccbe1^+/+^* (**G**) and *ccbe1^-/-^* (**H**) embryos carrying the *Tg(kdrl:EGFP)* transgene, showing severely impaired vascular formation at the pineal gland-dCP interface in the *ccbe1^-/-^* embryo compared with the well-formed vascular network in its control sibling. Cyan asterisks indicate sites of disrupted vascular formation. (**I, J**) Dorsal views of 48 hpf *ccbe1^+/+^* (**I**) and *ccbe1^-/-^* (**J**) embryos carrying the *Tg(kdrl:EGFP)* transgene, showing substantial recovery of vascular formation at the pineal gland-dCP interface in the *ccbe1^-/-^*embryo. (**K, L**) Quantification of vascular formation at the pineal gland-dCP interface in the indicated genotypes at 32 (**K**) and 48 (**L**) hpf (numbers of animals examined per genotype are listed in the panel). (**M, N**) Lateral views of 32 hpf *ccbe1^+/+^* (**M**) and *ccbe1^-/-^* (**N**) trunk vasculature visualized by the *Tg(kdrl:EGFP)* transgene, showing no obvious delay in trunk intersegmental artery (ISVa) development in the *ccbe1^-/-^* embryo (**N**). (O) Quantification of trunk ISVa development in the indicated genotypes at 32 hpf. (P) Experimental setup for the microinjection experiments shown in (**Q**-**X**). (**Q, R**) Dorsal views of 32 hpf Cas9-injected control (**Q**) and *ccbe1* crispant (**R**) embryos carrying the *Tg(tp1:Venus-PEST);Tg(kdrl:ras-mCherry)* reporters, showing severely impaired vascular formation at the pineal gland-dCP interface (**cyan asterisks**) in the *ccbe1* crispant (**R**), while pineal gland formation visualized by the *Tg(tp1:Venus-PEST)* transgene is largely preserved. (**S, T**) Dorsal views of 32 hpf Cas9-injected control (**S**) and *ccbe1* crispant (**T**) embryos carrying the heterozygous *ccbe1* knock-in allele with *Tg(UAS:EGFP);Tg(kdrl:ras-mCherry)* reporters, showing severely impaired vascular formation at the pineal gland-dCP interface (**cyan asterisks**) in the *ccbe1* crispant (**T**), while the formation of *ccbe1*-positive pineal gland cells is largely preserved. (**U**) Quantification of vascular formation at the pineal gland-dCP interface in 32 hpf Cas9-injected controls and *ccbe1* crispants. (**V, W**) Dorsal brightfield and fluorescent merged images of 32 hpf Cas9-injected control (**V**) and *ccbe1* crispant (**W**) embryos, showing the pineal gland visualized by the *Tg(tp1:Venus-PEST)* transgene. Pineal gland tissue size (outlined by dashed lines) remains largely preserved in *ccbe1* crispants. (X) Quantification of pineal gland tissue size in 32 hpf Cas9-injected controls and *ccbe1* crispants. Values represent mean ± SEM. Statistical significance was assessed by two-tailed Student’s *t*-test (NS: not significant). (Y) Summary of pineal gland and vascular development in 32 hpf control and *ccbe1* crispant embryos, showing largely preserved pineal gland development despite severely delayed vascularization in this brain region. In panels (**K, L, O**, and **U**), \*\*\*\**P* < 0.0001; Fisher’s exact test. NS: not significant. Scale bars: 50 µm in **A**-**C**; **H** for **G, H**; **J** for **I, J**; **N** for **M, N**; **R** for **Q, R**; **T** for **S, T**; **W** for **V, W**; 500 µm in **F** for **E, F**.

In *ccbe1* mutants, this stereotyped vascularization process was substantially delayed. At 32 hpf, the vascular pattern in *ccbe1* mutants closely resembled that observed in WT embryos approximately 10 hours earlier, indicating a marked delay in vascular development at the pineal gland-dCP interface (Fig. 2G, H, K). Intriguingly, despite this pronounced early defect, vascularization of this region largely recovered by 48 hpf (Fig. 2I, J, L). Thus, Ccbe1 is required for timely vascularization at the pineal gland-dCP interface. Using this phenotype as a readout, functional characterization of the *ccbe1* knock-in reporter revealed that the homozygous allele retains partial Ccbe1 function and behaves as a hypomorph (Fig. S1).

We next asked whether the delayed cranial vascularization in *ccbe1* mutants reflected a broader delay in embryonic angiogenesis. To address this possibility, we examined intersegmental vessel (ISV) formation in the trunk, where angiogenic sprouts emerge from the dorsal aorta and extend dorsally during early embryogenesis. In contrast to the pronounced delay at the pineal gland-dCP interface, trunk ISV formation proceeded with comparable timing in WT and *ccbe1* mutant embryos (Fig. 2M-O), consistent with their comparable overall embryonic growth (Fig. 2E, F). These findings indicate that loss of *ccbe1* does not cause a general delay in developmental angiogenesis or embryonic growth, but rather markedly delays vascularization of the pineal gland-dCP interface.

### Genetic loss of *ccbe1* causes severe vascular defects while largely preserving pineal gland development

Because impaired vascularization could potentially arise secondary to defects in pineal gland development, we next examined pineal gland morphology and quantified its size at 32 hpf. To this end, we employed a CRISPR/Cas9-based approach to generate F0 *ccbe1* knockout embryos (crispants) through injections of three distinct *ccbe1*-targeted dual-guide ribonucleoprotein (dgRNP) complexes at the one-cell stage (Fig. 2P). Notably, *ccbe1* crispants recapitulated the pronounced delay in vascularization at the pineal gland-dCP interface observed in stable *ccbe1* mutants (Fig. 2Q-U), validating this approach for effectively disrupting *ccbe1* function.

Leveraging this approach, we first visualized the pineal gland using the *Tg(tp1:Venus-PEST)* reporter together with DIC imaging. Comparison of Cas9-injected controls and *ccbe1* crispants revealed comparable overall pineal morphology despite the pronounced vascular defects in *ccbe1* crispants (Fig. 2Q, R). Consistent with this observation, we observed no significant difference in pineal gland size between the two groups (Fig. 2V-X). Importantly, despite its prominent expression, *ccbe1*-positive cells retained their organization within the pineal gland following *ccbe1* disruption (Fig. 2S, T). Combined with the severe vascular defects, these observations support a cell non-autonomous role for Ccbe1 in vascular development in this region.

Together, these findings indicate that *ccbe1* disruption causes a severe vascularization defect while gross pineal gland development is largely preserved (Fig. 2Y), suggesting that Ccbe1 is not essential for the early formation of the pineal gland.

### Vegfa signaling cooperates with the Ccbe1/Vegfc pathway during vascularization of the pineal gland-dCP interface

Although *ccbe1* loss caused a pronounced delay in vascularization at the pineal gland-dCP interface, vascular development in this region substantially recovered by 48 hpf. Given our recent evidence for functional interplay between Vegfc and Vegfa signaling during vascular development (*12, 13*), we asked whether Vegfa signaling contributes to this recovery and cooperates with Ccbe1 during cerebrovascular development. Zebrafish possess two Vegfa paralogs, Vegfaa and Vegfab. We therefore examined vascular development following combined genetic loss of *ccbe1* and either *vegfaa* or *vegfab*.

Compared with loss of *ccbe1* alone, we observed that combined loss of *ccbe1* and either *vegfa* paralog markedly exacerbated cerebrovascular defects. Analysis of *ccbe1;vegfaa* double mutants revealed a genetic interaction between *ccbe1* and *vegfaa* in overall vascularization at the pineal gland-dCP interface, leading to severely impaired formation of the prosencephalic artery (PrA), pineal gland vessel (PGV), and anterior cerebral vein (ACeV) (Fig. 3A-F). In contrast, *ccbe1;vegfab* double mutants exhibited more spatially restricted vascular defects, characterized by severely impaired PrA formation and milder deficits in PGV formation (Fig. 3I-N). These defects were substantially more pronounced than those observed in either corresponding single mutant (Fig. 3N), demonstrating a genetic interaction between *ccbe1* and *vegfab*. Importantly, these vascularization defects at the pineal gland-dCP interface persisted through at least 10 days post fertilization (dpf) (Fig. S2A-G). Thus, these findings reveal distinct, yet critical, genetic interactions of *ccbe1* with the two *vegfa* paralogs during vascularization at the pineal gland-dCP interface.

**Fig. 3.**
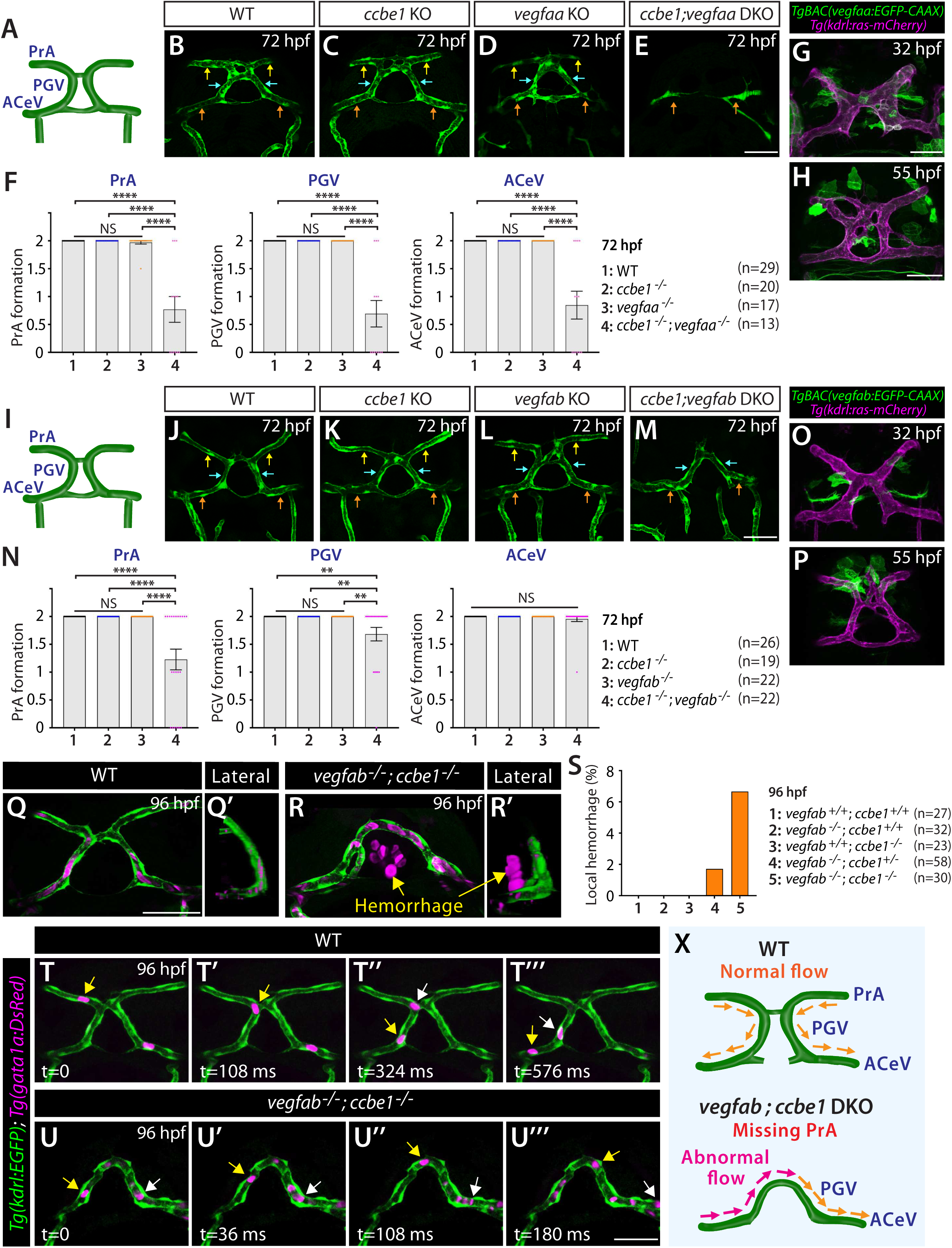
Combined loss of ccbe1 and Vegfa paralogs enhances angiogenic defects at the pineal gland-dCP interface and induces local hemorrhage and abnormal blood flow. (A) Schematic dorsal view of the vascular anatomy at the pineal gland-dCP interface at 72 hpf. (**B**-**E**) Dorsal views of the cranial vasculature at the pineal gland-dCP interface in 72 hpf WT (**B**), *ccbe1^-/-^* (**C**), *vegfaa^-/-^* (**D**), and *ccbe1^-/-^;vegfaa^-/-^* (**E**) larvae visualized by the *Tg(kdrl:*EGFP) transgene. Yellow arrows point to the PrA, blue arrows to the PGV, and orange arrows to the ACeV. (F) Quantification of PrA, PGV, and ACeV formation in the indicated genotypes at 72 hpf (numbers of animals examined per genotype are listed in the panel). *ccbe1* and *vegfaa* genetically interact in PrA, PGV, and ACeV formation. (**G, H**) Dorsal views of 32 and 55 hpf *TgBAC(vegfaa:gal4ff);Tg(UAS:EGFP-CAAX);Tg(kdrl:ras-mCherry)* embryonic heads, showing *TgBAC(vegfaa:gal4ff)*-driven EGFP expression in endothelial cells and non-endothelial cells associated with blood vessels at the pineal gland-dCP interface. (**I**) Schematic dorsal view of the vascular anatomy at the pineal gland-dCP interface at 72 hpf. (**J**-**M**) Dorsal views of the cranial vasculature at the pineal gland-dCP interface in 72 hpf WT (**J**), *ccbe1^-/-^* (**K**), *vegfab^-/-^* (**L**), and *ccbe1^-/-^;vegfab^-/-^* (**M**) larvae visualized by the *Tg(kdrl:*EGFP) transgene. Yellow arrows point to the PrA, blue arrows to the PGV, and orange arrows to the ACeV. The *ccbe1^-/-^;vegfab^-/-^*larva failed to form bilateral PrAs (**M**). (**N**) Quantification of PrA, PGV, and ACeV formation in the indicated genotypes at 72 hpf (numbers of animals examined per genotype are listed in the panel). *ccbe1* and *vegfab* genetically interact in PrA and PGV formation, with a particularly notable increase in PrA formation defects. (**O, P**) Dorsal views of 32 and 55 hpf *TgBAC(vegfab:gal4ff);Tg(UAS:EGFP-CAAX);Tg(kdrl:ras-mCherry)* embryonic heads, showing *TgBAC(vegfab:gal4ff)*-driven EGFP expression in non-endothelial cells associated with blood vessels at the pineal gland-dCP interface. At 55 hpf, these EGFP-positive cells are prominently localized near the PrA (**P**). (**Q**-**R’**) Dorsal (**Q, R**) and lateral (**Q’, R’**) views of the cranial vasculature and red blood cells at the pineal gland-dCP interface in 96 hpf WT (**Q, Q’**) and *ccbe1^-/-^;vegfab^-/-^* (**R, R’**) larvae visualized by the *Tg(kdrl:*EGFP) and *Tg(gata1a:DsRed)* reporters. Bilateral PrA loss and local hemorrhage at the pineal gland-dCP interface were observed in the *ccbe1^-/-^;vegfab^-/-^* (**R, R’**) larva. (**S**) Quantification of local hemorrhage at the pineal gland-dCP interface in the indicated genotypes at 96 hpf. (**T**-**U’’’**) Time-lapse tracking of red blood cell movement in 96 hpf WT (**T**-**T’’’**) and *ccbe1^-/-^;vegfab^-/-^* (**U**-**U’’’**) larvae visualized by the *Tg(kdrl:*EGFP) and *Tg(gata1a:DsRed)* reporters. Yellow and white arrows indicate the positions of tracked red blood cells. While the WT larva exhibits directional red blood cell movement from the PrA to the ACeV through the PGV, the *ccbe1^-/-^;vegfab^-/-^* larva lacking bilateral PrAs exhibits abnormal blood flow from one side of the ACeV to the other. (**X**) Schematic diagrams illustrating the observed blood flow directions in 96 hpf WT and *ccbe1^-/-^;vegfab^-/-^* larvae shown in (**T**-**U’’’**). In panels (**F** and **N**), each data point represents an individual animal’s vessel formation score, and values represent mean ± SEM (\*\**P* < 0.01 and \*\*\*\**P* < 0.0001; one-way ANOVA followed by Tukey’s multiple-comparison test; NS: not significant). Scale bars: 50 µm in **E** for **B**-**E**; **G** for **G, O**; **H** for **H, P**; **M** for **J**-**M**; **Q** for **Q, R**; **U’’’** for **T**-**U’’’**.

The differential requirements of *vegfaa* and *vegfab* in regulating this vascularization process motivated us to examine their expression patterns. To this end, we used their BAC transgenic reporters for *vegfaa* and *vegfab*, which exhibited spatially restricted expression domains within the developing brain, including regions in close proximity to the pineal gland-dCP vasculature (Fig. 3G, H, O, P). Specifically, the *vegfaa* BAC reporter exhibited both EC and non-EC expression at this interface at 32 hpf (Fig. 3G). These expression patterns largely persisted through 55 hpf, although EC reporter expression appeared reduced at this stage (Fig. 3H). In contrast, the *vegfab* BAC reporter initially displayed relatively sparse non-EC expression at this interface at 32 hpf (Fig. 3O), which became more localized to the region in close proximity to the PrAs by 55 hpf (Fig. 3P). These distinct expression patterns are consistent with different spatial contributions of the two Vegfa paralogs to vascularization at this interface.

These findings suggested that Vegfa signaling functions cooperatively with Ccbe1 during vascularization at the pineal gland-dCP interface. To determine whether these genetic requirements were specific to Vegfa signaling, we next examined *ccbe1;vegfc* double mutants (Fig. S3A). In contrast to the markedly enhanced vascular defects observed following combined loss of *ccbe1* and either *vegfa* paralog at this interface, combined loss of *ccbe1* and *vegfc* did not substantially exacerbate vascular defects compared with the corresponding single mutants (Fig. S3B, C, F). These observations are consistent with Ccbe1 and Vegfc functioning within the same developmental pathway, as previously demonstrated in various biological contexts (*24, 25, 27*).

### Vascular defects at the pineal gland-dCP interface are associated with local hemorrhage and altered blood flow

We next asked whether developmental vascularization defects at the pineal gland-dCP interface have physiological consequences. To address this question in live larvae, we used the *Tg(gata1a:DsRed)* reporter, which labels erythrocytes and enables visualization of blood flow (*39*). During these analyses, we occasionally observed substantial accumulation of erythrocytes in tissues surrounding the pineal gland, indicative of local hemorrhage (Fig. 3Q-R’). We therefore quantified the incidence of this phenotype and observed a higher incidence of local hemorrhage in *ccbe1;vegfab* double mutants than in the corresponding single mutants, although this difference did not reach statistical significance (Fig. 3S). Notably, hemorrhage was primarily observed in larvae exhibiting incomplete or abnormal vessel formation rather than complete vessel loss. Because *ccbe1;vegfab* double mutants frequently exhibited complete loss of specific vessels in this region, the absence of these vessels may partially account for the relatively low overall incidence of hemorrhage.

To further determine how these vascular defects affect local circulation, we examined blood flow through the pineal gland-dCP interface using the *Tg(gata1a:DsRed)* reporter. In WT larvae, blood flowed from the PrA through the PGV toward the ACeV at this interface (Fig. 3T-T’’’; Movie S1). In contrast, *ccbe1;vegfab* double mutants lacking the bilateral PrAs exhibited a markedly altered flow pattern, in which blood entered the PGV from the ACeV on one side and flowed toward the contralateral ACeV (Fig. 3U-U’’’; Movie S2). Thus, loss of the bilateral PrAs redirects blood flow through the remaining vascular connections, substantially altering local circulation at the pineal gland-dCP interface.

Together, the occurrence of local hemorrhage and altered blood flow in *ccbe1;vegfab* double mutants demonstrate that developmental vascularization defects at the pineal gland-dCP interface are associated with physiological alterations in local cerebrovascular function.

### Ccbe1 cooperates with Vegfab to promote fenestrated vessel formation in the hindbrain mCP

We next asked whether the requirement for Ccbe1 in brain vascularization extends beyond the pineal gland-dCP interface and represents a broader mechanism contributing to the development of fenestrated cerebrovascular beds. To address this question, we examined vascularization of the mCP, another specialized fenestrated vascular bed located in the hindbrain. This investigation was further motivated by *ccbe1* knock-in reporter and endogenous *ccbe1* expression detected near the dorsal midline junction (DMJ) (Fig. 4A, B), where a major vessel contributing to mCP vascularization originates.

**Fig. 4.**
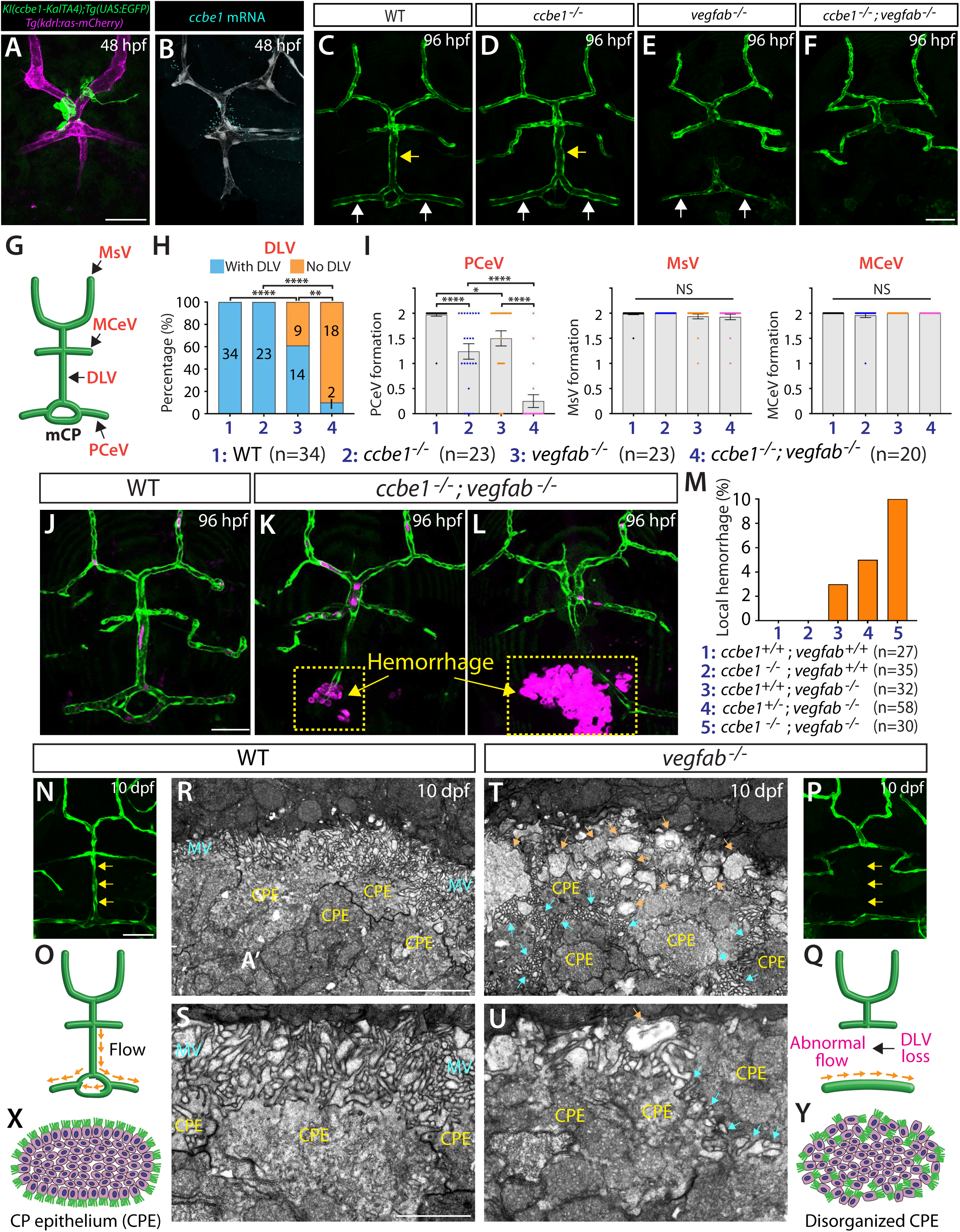
Ccbe1 cooperates with Vegfa signaling to promote fenestrated vessel formation in the hindbrain myelencephalic CP (mCP), with impaired mCP vascularization associated with local hemorrhage and epithelial disorganization. (**A, B**) Side-by-side comparison of a 48 hpf homozygous *ccbe1* knock-in embryo carrying the *Tg(UAS:EGFP-CAAX);Tg(kdrl:ras-mCherry)* reporters (**A**) and *ccbe1* mRNA expression (**B**), showing consistent positive signals at the dorsal midline junction (DMJ), where DLV sprouting initiates. (**C**-**F**) Dorsal views of the cranial vasculature near the mCP in 96 hpf WT (**C**), *ccbe1^-/-^* (**D**), *vegfab^-/-^* (**E**), and *ccbe1^-/-^;vegfab^-/-^* (**F**) larvae visualized by the *Tg(kdrl:*EGFP) transgene, showing loss of the DLV in *vegfab^-/-^* larva and complete loss of the DLV and bilateral PCeVs in *ccbe1^-/-^;vegfab^-/-^* larva. Yellow arrows point to the DLV and white arrows to the PCeV. (G) Schematic dorsal view of the vascular anatomy near the mCP at 96 hpf, indicating the blood vessels examined in this brain region. **MsV**: mesencephalic vein, **MCeV**: middle cerebral vein, **DLV**: dorsal longitudinal vein, **PCeV**: posterior cerebral vein. (H) Percentage of 96 hpf larvae of the indicated genotypes with or without the DLV (numbers of animals examined per genotype are listed in the panel), showing a strong genetic interaction between *ccbe1* and *vegfab* in DLV formation. \*\**P* < 0.01 and \*\*\*\**P* < 0.0001; Fisher’s exact test. (I) Quantification of PCeV, MsV, and MCeV formation in the indicated genotypes at 96 hpf, showing a strong genetic interaction between *ccbe1* and *vegfab* in PCeV formation, whereas MsV and MCeV formation remains unaffected in *ccbe1^-/-^;vegfab^-/-^*larvae. Each data point represents an individual animal’s vessel formation score, and values represent mean ± SEM (\**P* < 0.05 and \*\*\*\**P* < 0.0001; one-way ANOVA followed by Tukey’s multiple-comparison test; NS: not significant). (**J**-**L**) Dorsal views of the cranial vasculature and red blood cells near the mCP in 96 hpf WT (**J**) and *ccbe1^-/-^;vegfab^-/-^* (**K, L**) larvae visualized by the *Tg(kdrl:*EGFP) and *Tg(gata1a:DsRed)* reporters. Incomplete mCP vascularization is associated with local hemorrhage (arrows) near the mCP in the *ccbe1^-/-^;vegfab^-/-^* larvae (**K, L**). (**M**) Quantification of local hemorrhage near the mCP in the indicated genotypes at 96 hpf. (**N, P**) Dorsal views of the cranial vasculature near the mCP in 10 dpf WT (**N**) and *vegfab^-/-^* (**P**) larvae visualized by the *Tg(kdrl:*EGFP) transgene, showing loss of the DLV in *vegfab^-/-^* larva. (**O, Q**) Schematic diagrams illustrating the cranial vasculature near the mCP and blood flow directions in 10 dpf WT (**O**) and *vegfab^-/-^* (**Q**) larvae shown in (**N, P**). (**R**-**U**) Transmission electron microscopy (TEM) images of the mCP in 10 dpf WT (**R, S**) and DLV-lacking *vegfab^-/-^* (**T, U**) larvae. Magnified images are shown in (**S, U**). The WT larva exhibits a well-organized epithelial cell layer characterized by centrally located cell bodies and microvilli extending toward the surrounding tissues (**R, S**). In contrast, the DLV-lacking *vegfab^-/-^* larva exhibits disrupted epithelial cell organization, including apparently misoriented microvilli (cyan arrows in **T, U**) and ectopic swollen-like cellular structures (orange arrows in **T, U**). CPE: choroid plexus epithelial cell; MV: microvilli. (**X, Y**) Schematic diagrams illustrating mCP epithelial cell organization in 10 dpf WT (**X**) and *vegfab^-/-^* (**Y**) larvae based on the TEM images shown in (**R**-**U**). Scale bars: 50 µm in **A** for **A, B**; **F** for **C**-**F**; **J** for **J**-**L**; **N** for **N, P**; 5 µm in **R** for **R, T**; 2 µm in **S** for **S, U**.

The developing mCP is vascularized by the dorsal longitudinal vessel (DLV) and bilateral posterior cerebral veins (PCeVs), which form a stereotyped vascular network closely associated with the mCP epithelium (*40*) and exhibit molecular signatures characteristic of fenestrated vessels (*13, 14*). Given the prominent requirement for Vegfab in mCP vascularization (*13*), we focused on examining *ccbe1* mutants alone and in combination with *vegfab* mutants. At 96 hpf, *ccbe1* single mutants exhibited largely normal mCP vascularization, with no apparent defects in either DLV or PCeV formation (Fig. 4C, D). In contrast, combined loss of *ccbe1* and *vegfab* caused severe defects in mCP vascularization, affecting formation of both the DLV and PCeVs (Fig. 4F) and markedly enhancing the DLV-loss phenotype observed in *vegfab* single mutants (Fig. 4E-H). Importantly, while these mCP vascularization defects persisted through 10 dpf (Fig. S2H-O), formation of the neighboring mesencephalic vein (MsV) and middle cerebral vein (MCeV), which eventually acquire BBB features, was largely unaffected in the double mutants (Fig. 4I). These observations indicate that the vascular defects were spatially restricted rather than reflecting a general impairment of hindbrain angiogenesis.

Together, these findings reveal a strong genetic interaction between *ccbe1* and *vegfab* in this vascularization context and demonstrate a further cooperative requirement for Ccbe1 and Vegfa signaling in the formation of the fenestrated cerebrovascular network.

### Defective mCP vascularization is associated with structural disorganization of the CP epithelium

We next asked whether these severe mCP vascularization defects in *ccbe1;vegfab* double mutants were associated with compromised vascular integrity. Using the *Tg(gata1a:DsRed)* reporter to visualize erythrocytes, we occasionally observed substantial accumulation of erythrocytes in tissues surrounding the mCP, indicative of local hemorrhage (Fig. 4J-L). Quantification revealed a progressively higher incidence of local hemorrhage with increasing loss of *ccbe1* alleles in the *vegfab* mutant background, from *vegfab^-/-^*larvae to *vegfab^-/-^;ccbe1^+/-^* and *vegfab^-/-^;ccbe1^-/-^*animals (Fig. 4M). Notably, hemorrhage was predominantly associated with partial or abnormal vessel formation rather than complete loss of the affected vessels. Because complete loss of specific mCP vessels occurred frequently in *ccbe1;vegfab* double mutants, the absence of these vascular conduits may partially account for the relatively low overall incidence of hemorrhage in these animals despite the severity of the vascular phenotype.

The CP contributes to brain homeostasis through cerebrospinal fluid production, waste clearance, and formation of the blood-CSF barrier (*41*). Given the close association between fenestrated vessels and the CP epithelium, we next asked whether defective mCP vascularization was accompanied by alterations in epithelial organization. To distinguish abnormalities associated with altered vascular architecture from those resulting from complete loss of blood circulation, we analyzed larvae in which the DLV was absent while the bilateral PCeVs remained intact. This vascular pattern maintained blood circulation around the mCP while altering its local flow pattern (Fig. 4N-Q).

Ultrastructural analysis revealed pronounced abnormalities in the organization of the mCP epithelium (Fig. 4R-U; Fig. S4). In WT larvae, CP epithelial cells formed an organized epithelial layer with microvilli projecting from their apical surface into the ventricular space. In larvae with defective mCP vascularization, epithelial organization was disrupted, with microvilli extending into aberrant locations and ectopic swollen-like enlarged subcellular structures that were not apparent in WT larvae (Fig. 4R-Y; Fig. S4). Thus, these observations suggest that impaired formation of the mCP-associated vascular network is associated with epithelial disorganization.

Together, these findings reveal an association between proper mCP vascularization and the structural organization of the CP epithelium, indicating that local vascular morphogenesis may contribute to epithelial integrity at the blood-CSF barrier.

### *ccbe1* and *vegfc* mutants closely phenocopy brain region-specific angiogenic defects

Ccbe1-dependent proteolytic activation of Vegfc is crucial for lymphangiogenesis, providing spatiotemporal control of Vegfc signaling during lymphatic development (*23, 25*). We therefore asked whether Ccbe1 and Vegfc function similarly during brain angiogenesis. To address this question, we systematically compared cerebrovascular phenotypes between *ccbe1* and *vegfc* mutants, with particular attention to brain regions exhibiting prominent Ccbe1-dependent vascularization defects.

Remarkably, we found that *vegfc* single mutants exhibited highly similar region-specific angiogenic defects to *ccbe1* single mutants, including impaired vascularization at the pineal gland-dCP interface (Fig. 5C-E). In support of these phenotypic similarities, the *vegfc* BAC reporter showed robust EC and non-EC expression at this interface in close proximity to *ccbe1* knock-in reporter-positive cells in the pineal gland at 26 hpf (Fig. 5A, B).

**Fig. 5.**
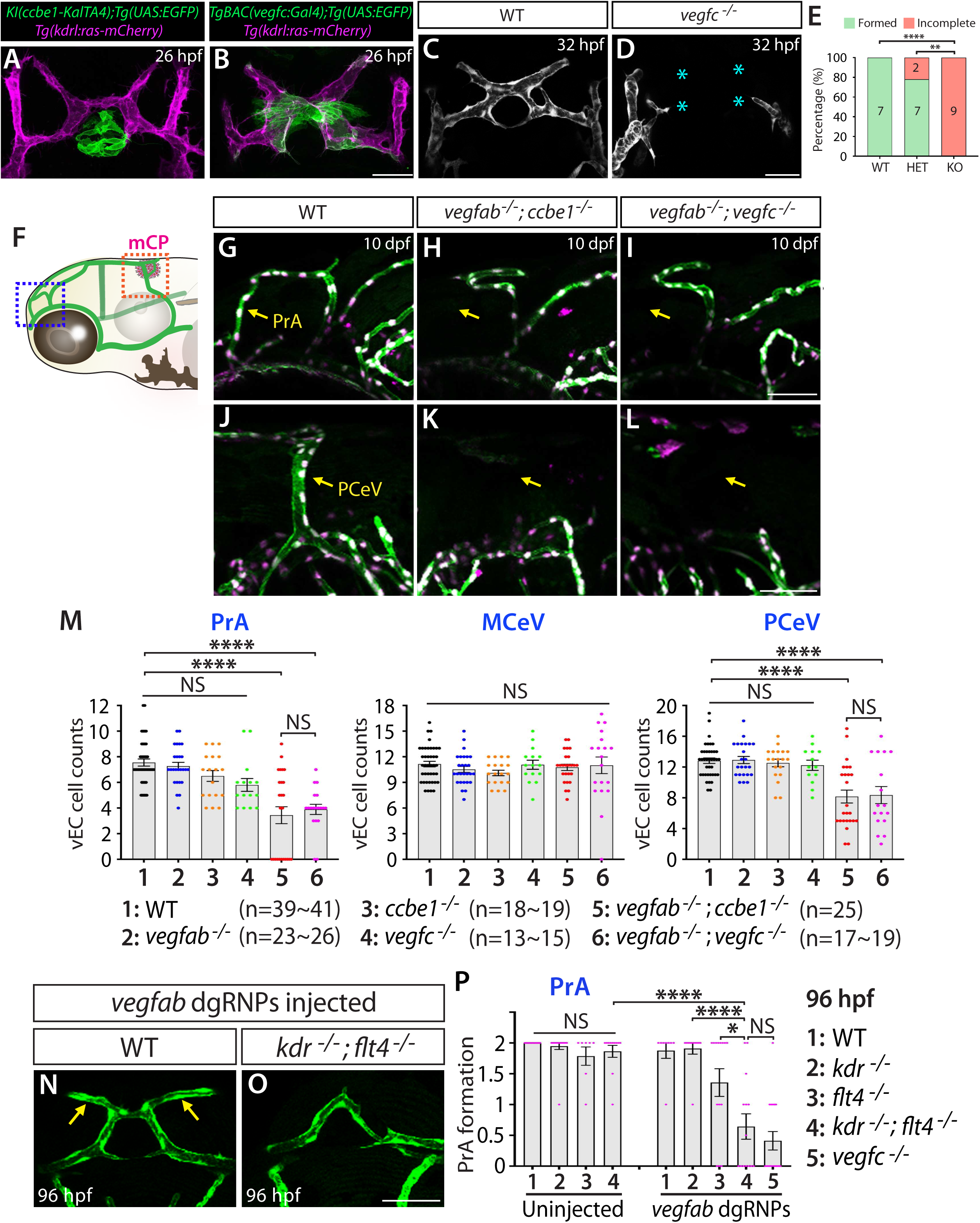
*ccbe1* and *vegfc* mutants closely phenocopy brain region-specific angiogenic defects. (**A, B**) Dorsal views of 26 hpf *ccbe1* knock-in heterozygous (**A**) and *TgBAC(vegfc:gal4ff)* (**B**) embryos carrying the *Tg(UAS:EGFP-CAAX);Tg(kdrl:ras-mCherry)* reporters, showing prominent *ccbe1* knock-in reporter expression in the developing pineal gland and *vegfc* BAC reporter expression around this brain region, including endothelial cells and non-endothelial cell populations. (**C, D**) Dorsal views of 32 hpf *vegfc^+/+^* (**C**) and *vegfc^-/-^*(**D**) embryos carrying the *Tg(kdrl:EGFP)* transgene, showing severely impaired vascular formation at the pineal gland-dCP interface in the *vegfc^-/-^*embryo compared with the well-formed vascular network in its control sibling. Cyan asterisks indicate sites of disrupted vascular formation. (E) Quantification of vascular formation at the pineal gland-dCP interface in the indicated genotypes at 32 hpf (numbers of animals examined per genotype are listed in the panel). \*\**P* < 0.01 and \*\*\*\**P* < 0.0001; Fisher’s exact test. (F) Simplified schematic diagram indicating the two lateral locations used to assess vascular phenotypes in (**G**-**M**). (**G**-**L**) Lateral views of the cranial vasculature near the pineal gland-dCP interface (**G**-**I**) and mCP region (**J**-**L**) in 10 dpf WT (**G, J**), *vegfab^-/-^;ccbe1^-/-^* (**H, K**), and *vegfab^-/-^;vegfc^-/-^* (**I, L**) larvae visualized by the *Tg(kdrl:EGFP)* and *Tg(kdrl:NLS-mCherry)* reporters, showing loss of the PrA (arrows in **H, I**) and PCeV (arrows in **K, L**) in both double mutants. (**M**) Quantification of endothelial cell numbers constituting the PrA, MCeV, and PCeV at 10 dpf, showing considerable and similar reductions in PrA and PCeV endothelial cell numbers in both *vegfab^-/-^;ccbe1^-/-^* and *vegfab^-/-^;vegfc^-/-^* larvae, whereas MCeV endothelial cell numbers remain comparable to WT. (**N, O**) Dorsal views of the cranial vasculature at the pineal gland-dCP interface in 96 hpf WT (**N**) and *kdr^-/-^;flt4^-/-^*(**O**) larvae injected with *vegfab* dgRNPs and visualized by the *Tg(kdrl:EGFP)* transgene, showing bilateral loss of the PrAs in *vegfab* dgRNPs-injected *kdr^-/-^;flt4^-/-^* larvae. (**P**) Quantification of PrA formation under the indicated conditions at 96 hpf, showing strong genetic interactions between *kdr;flt4* and *vegfab* in PrA formation and phenotypic similarity to the combined loss of *vegfc* and *vegfab*. In panel (**M**), each data point represents an individual animal’s EC cell count; in panel (**P**), each data point represents an individual animal’s PrA formation score. Values represent mean ± SEM (\**P* < 0.05 and \*\*\*\**P* < 0.0001; one-way ANOVA followed by Tukey’s multiple-comparison test; NS: not significant). Scale bars: 50 µm in **B** for **A, B**; **D** for **C, D**; **I** for **G**-**I**; **L** for **J**-**L**; **O** for **N, O**.

To compare cerebrovascular phenotypic similarities between *ccbe1* and *vegfc* under reduced Vegfa conditions, we used *Tg(kdrl:EGFP);Tg(kdrl-NLS-mCherry)* reporter lines, which visualize EC cell bodies and nuclei, respectively, enabling precise quantification of vascular EC numbers. Using this approach, quantitative analyses revealed closely corresponding reductions in these cell numbers between *ccbe1;vegfab and vegfc;vegfab* mutants in vessels associated with both the pineal gland-dCP interface and mCP, including the PrA and PCeV, respectively (Fig. 5F-M). In contrast, EC numbers remained largely unaffected in the MCeV, a vessel with BBB characteristics (Fig. 5M). Together with the vascular phenotypic similarities observed in the single mutants, these quantitative comparisons suggest a model in which Ccbe1 functions through the Vegfc pathway to promote region-specific brain angiogenesis. Our earlier observation that combined loss of *ccbe1* and *vegfc* did not further enhance vascular defects at either the pineal gland-dCP interface or mCP (Fig. S3) further supports this model.

### Flt4 and Vegfr2 receptors mediate Ccbe1/Vegfc-and Vegfab-dependent vascularization of the pineal gland-dCP interface

Vegfc-Vegfr3 signaling is essential for lymphatic development in both zebrafish and mice (*27, 28, 42, 43*). Previous biochemical studies in zebrafish further demonstrated that Vegfc can bind both Flt4 (Vegfr3) and Kdr, one of the two zebrafish Vegfr2 orthologs (*44*). We therefore asked whether Flt4 and Kdr mediate Ccbe1/Vegfc-dependent brain vascularization. To address this question, we analyzed the *flt4^um131^*and/or *kdr^bns32^* mutant alleles individually and in combination, with or without simultaneous CRISPR-mediated disruption of *vegfab*.

In the absence of *vegfab* disruption, *flt4;kdr* double mutant larvae exhibited no substantial defects in vascularization at the pineal gland-dCP interface (Fig. 5P). In contrast, simultaneous disruption of *vegfab* revealed a strong requirement for these receptors: *vegfab*-targeted *flt4;kdr* double mutants exhibited pronounced defects in PrA formation, whereas the corresponding single receptor mutants displayed substantially weaker phenotypes (Fig. 5N-P). These findings indicate that Flt4 and Kdr function cooperatively to promote PrA formation under reduced Vegfab signaling. Combined with our recent study showing that Flt4 and Kdr cooperatively regulate mCP vascularization under reduced Vegfa signaling (*14*), and given the potential contribution of Kdrl as another zebrafish Vegfr2 ortholog, these findings support a model in which Flt4 together with Vegfr2 receptors mediates the cooperative effects of Ccbe1/Vegfc and Vegfab signaling during brain vascularization.

### Expression and genetic analyses indicate brain region-specific requirements for Adamts3 and Adamts14 during vascularization

The secreted protein CCBE1 acts as a critical cofactor that facilitates proteolytic activation of VEGF-C by ADAMTS metalloproteinases (*22–26*). In particular, Adamts3 and Adamts14 function together with Ccbe1 as established proteolytic activators of Vegfc in multiple developmental and pathological contexts, including lymphangiogenesis (*23–26, 45, 46*). Genetic and biochemical studies further indicate that this proteolytic mechanism is conserved between mice and zebrafish (*24, 25, 32*). We next sought to define Vegfc-activating proteases that contribute to regional brain vascularization. This investigation was further motivated by the previously uncharacterized contributions and spatial requirements of Adamts3 and Adamts14 during brain vascularization, despite extensive characterization of the Vegfc-Ccbe1-Adamts3/14 signaling axis in lymphatic development.

To define their expression and vascular functions in the developing brain, we generated new knock-in reporter lines for *adamts3* and *adamts14* (Fig. 6A, B). Using a strategy similar to our *ccbe1* knock-in reporter, each line was generated by CRISPR/Cas9-mediated integration of an E2A-KalTA4 cassette into the endogenous locus (Fig. S5A, B). Crossing these lines to *Tg(UAS:EGFP-CAAX)* fish enabled membrane-localized fluorescent labeling of *adamts3-* and *adamts14*-expressing cells in live animals at single-cell resolution. Since individual loss of *adamts3* or *adamts14* does not produce overt cerebrovascular or trunk lymphatic defects (*32*), the extent of residual Adamts3 or Adamts14 function retained by the corresponding knock-in alleles was not rigorously assessed.

**Fig. 6.**
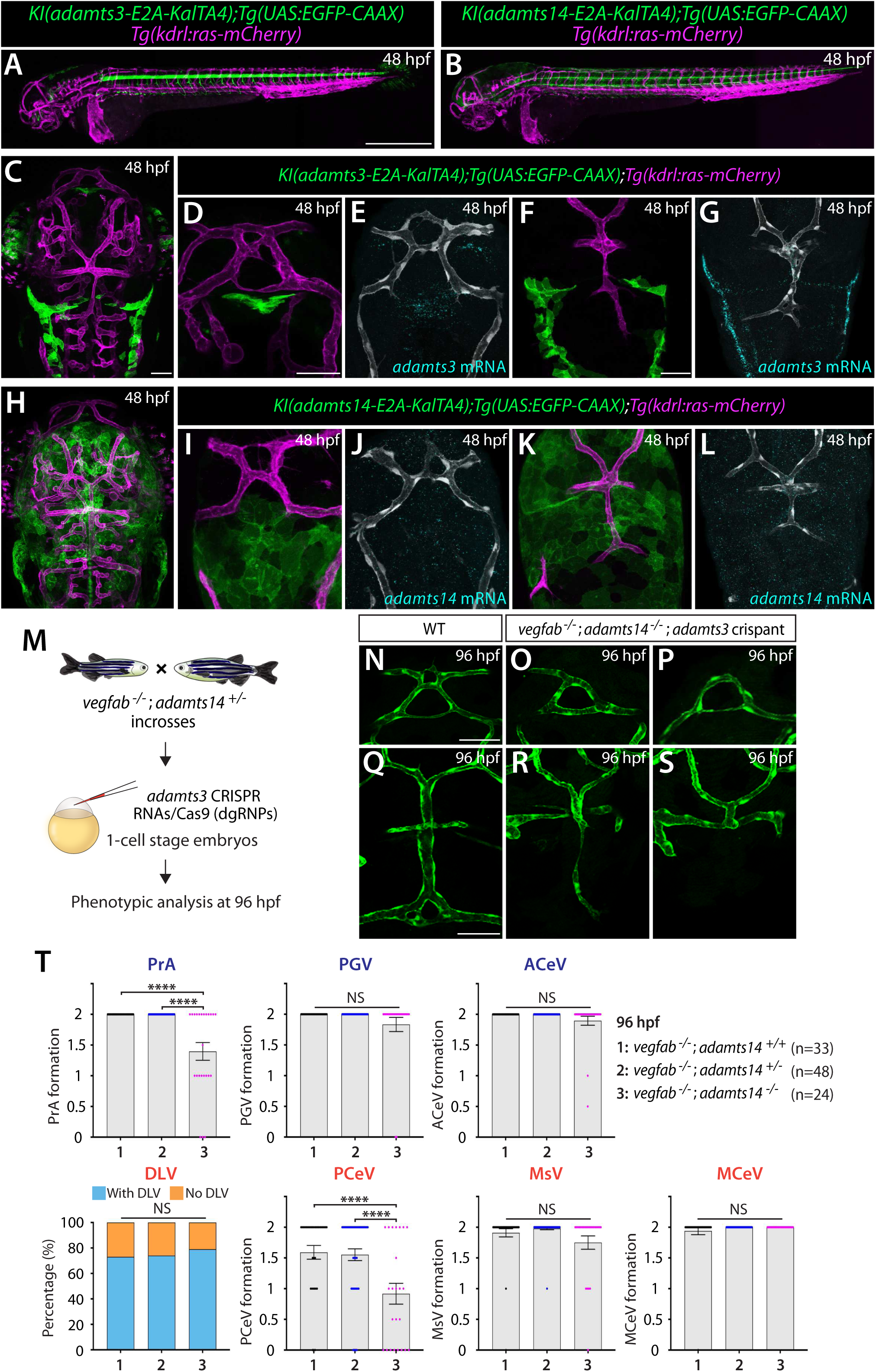
Expression and genetic analyses indicate brain region-specific requirements for Adamts3 and Adamts14 during vascularization. (**A, B**) Lateral views of 48 hpf *adamts3* (**A**) and *adamts14* (**B**) knock-in homozygous whole embryos carrying the *Tg(UAS:EGFP-CAAX);Tg(kdrl:ras-mCherry)* reporters. (**C**) Dorsal view of a 48 hpf *adamts3* knock-in homozygous embryonic head carrying the *Tg(UAS:EGFP-CAAX);Tg(kdrl:ras-mCherry)* reporters. (**D**-**G**) Side-by-side comparisons of 48 hpf heterozygous *adamts3* knock-in embryos carrying the *Tg(UAS:EGFP-CAAX);Tg(kdrl:ras-mCherry)* reporters (**D, F**) and *adamts3* mRNA expression (**E, G**), showing consistent signal patterns at the pineal gland-dCP interface and mCP regions. (**H**) Dorsal view of a 48 hpf *adamts14* knock-in homozygous embryonic head carrying the *Tg(UAS:EGFP-CAAX);Tg(kdrl:ras-mCherry)* reporters. (**I**-**L**) Side-by-side comparisons of 48 hpf heterozygous *adamts14* knock-in embryos carrying the *Tg(UAS:EGFP-CAAX);Tg(kdrl:ras-mCherry)* reporters (**I, K**) and *adamts14* mRNA expression (**J, L**), showing consistent signal patterns at the pineal gland-dCP interface and mCP regions. (**M**) Experimental setup for the microinjection experiments shown in (**N**-**T**). (**N**-**S**) Dorsal views of the cranial vasculature in 96 hpf WT (**N, Q**) and *vegfab^-/-^;adamts14^-/-^;adamts3* crispant (**O, P, R, S**) larvae visualized by the *Tg(kdrl:EGFP)* reporter, showing unilateral or bilateral loss of PrAs in mutants (**O, P**) and loss of bilateral PCeVs with or without the DLV (**R, S**). WT images in panels (**N**) and (**Q**) obtained from separate crosses are included as reference images. (**T**) Quantification of vascular formation at the pineal gland-dCP interface and mCP regions in the indicated genotypes at 96 hpf, showing significantly enhanced defects in PrA and PCeV formation in *vegfab^-/-^;adamts14^-/-^;adamts3* crispant larvae. Each data point represents an individual animal’s vessel formation score, and values represent mean ± SEM (\*\*\*\**P* < 0.0001; one-way ANOVA followed by Tukey’s multiple-comparison test; NS: not significant). For DLV quantification, statistical significance was assessed by Fisher’s exact test (NS: not significant). Scale bars: 500 µm in **A** for **A, B**; 50 µm in **C** for **C, H**; **D** for **D, E, I, J**; **F** for **F, G, K, L**; **N** for **N**-**P**; **Q** for **Q**-**S**.

Analysis of these reporters revealed distinct spatial patterns of *adamts3* and *adamts14* expression in the developing brain (Fig. 6C-L; Fig. S5C-F). In contrast to the prominent and highly localized *ccbe1* expression within the pineal gland (Fig. 1G, H), *adamts3* and *adamts14* reporter expression in this organ was comparatively subtle (Fig. S6). However, both reporters labeled cells in meningeal and perivascular regions closely associated with blood vessels at the pineal gland-dCP interface (Fig. 6D, E, I, J). Notably, *adamts3* exhibited more spatially restricted expression (Fig. 6D, E) compared with the broader distribution of *adamts14* expression (Fig. 6I, J). Both *adamts3*-and *adamts14*-expressing cells were also observed in proximity to developing vessels associated with the mCP, where *adamts3* expression was similarly more restricted (Fig. 6F, G), whereas *adamts14* expression was broadly distributed (Fig. 6K, L). Thus, the three components implicated in Vegfc proteolytic activation exhibit distinct but partially overlapping spatial distributions across the developing brain.

These expression patterns are consistent with previous studies identifying mesenchymal stromal cells, including *pdgfra*-positive fibroblasts, as major sources of secreted lymphangiogenic factors such as Vegfc and Ccbe1 during lymphangiogenesis (*32, 33*). Similarly, our recent study identified meningeal fibroblasts near the mCP as a source of multiple Vegf ligands, including Vegfab, Vegfc, and Vegfd (*13*). Together, these observations indicate that meningeal and perivascular regions provide important local sources of signaling components involved in Vegfc-dependent cerebrovascular development.

We next asked whether Adamts3 and Adamts14 are functionally required for vascularization of these specialized brain regions. Individual loss of *adamts3* or *adamts14* did not cause substantial vascularization defects in either WT or *vegfab* mutant backgrounds (Fig. S7, S8). We therefore tested whether Adamts3 and Adamts14 function cooperatively during brain vascularization. To simultaneously disrupt both genes, we incrossed *vegfab^-/-^;adamts14^+/-^* adults and subjected their progeny to CRISPR-mediated *adamts3* inactivation using three independent *adamts3*-targeting dgRNPs at the one-cell stage (Fig. 6M). In *vegfab^-/-^;adamts14^+/+^* larvae, *adamts3* disruption did not substantially affect vascularization at either the pineal gland-dCP interface or mCP (Fig. 6T). In contrast, *adamts3* disruption in *vegfab^-/-^;adamts14^-/-^* larvae significantly impaired PrA formation, revealing a requirement for combined Adamts3 and Adamts14 function in the *vegfab* mutant background (Fig. 6N-P).

A similar requirement for both proteases was observed during mCP vascularization. *vegfab^-/-^;adamts14^-/-^* larvae subjected to *adamts3* disruption exhibited impaired PCeV formation, whereas neighboring vessels remained largely unaffected (Fig. 6Q-T). These findings indicate that Adamts3 and Adamts14 together promote formation of the PrA and PCeV at the pineal gland-dCP interface and mCP, respectively. Interestingly, however, simultaneous disruption of *adamts3* and *adamts14* in the *vegfab* mutant background did not substantially impair DLV formation (Fig. 6T), suggesting distinct proteolytic requirements among individual cerebrovascular beds.

### Pineal gland-specific overexpression of activated Vegfc partially rescues vascularization defects caused by combined loss of *ccbe1* and *vegfab*

Our genetic analyses support a model in which Ccbe1 and Adamts3/14 function through the Vegfc signaling pathway during brain vascularization, raising the possibility that localized Ccbe1/Adamts-dependent proteolysis spatially controls Vegfc activity at the pineal gland-dCP interface. If Ccbe1 and Adamts3/14 promote vascularization primarily by enabling local Vegfc activation, pineal gland-specific expression of proteolytically activated Vegfc should bypass the requirement for Ccbe1. We therefore tested whether spatially restricted expression of activated Vegfc could restore vascularization following combined loss of *ccbe1* and *vegfab*.

To achieve spatial control of Vegfc activity, we generated a transgenic line expressing a previously validated activated form of zebrafish Vegfc (amino acids 97–221) (*44*) under the control of *aanat2* regulatory elements, thereby restricting Vegfc overexpression to the pineal gland. The transgene was designed to co-express mTagBFP2 and activated Vegfc separated by a P2A peptide, enabling identification of transgene-expressing pineal cells (Fig. 7A). To assess the effects of pineal gland-specific activated Vegfc expression, we simultaneously disrupted *ccbe1* and *vegfab* by subjecting *vegfab* mutants to CRISPR-mediated *ccbe1* inactivation (Fig. 7B), an approach that robustly recapitulated the PrA defects observed in stable *ccbe1;vegfab* double mutants.

**Fig. 7.**
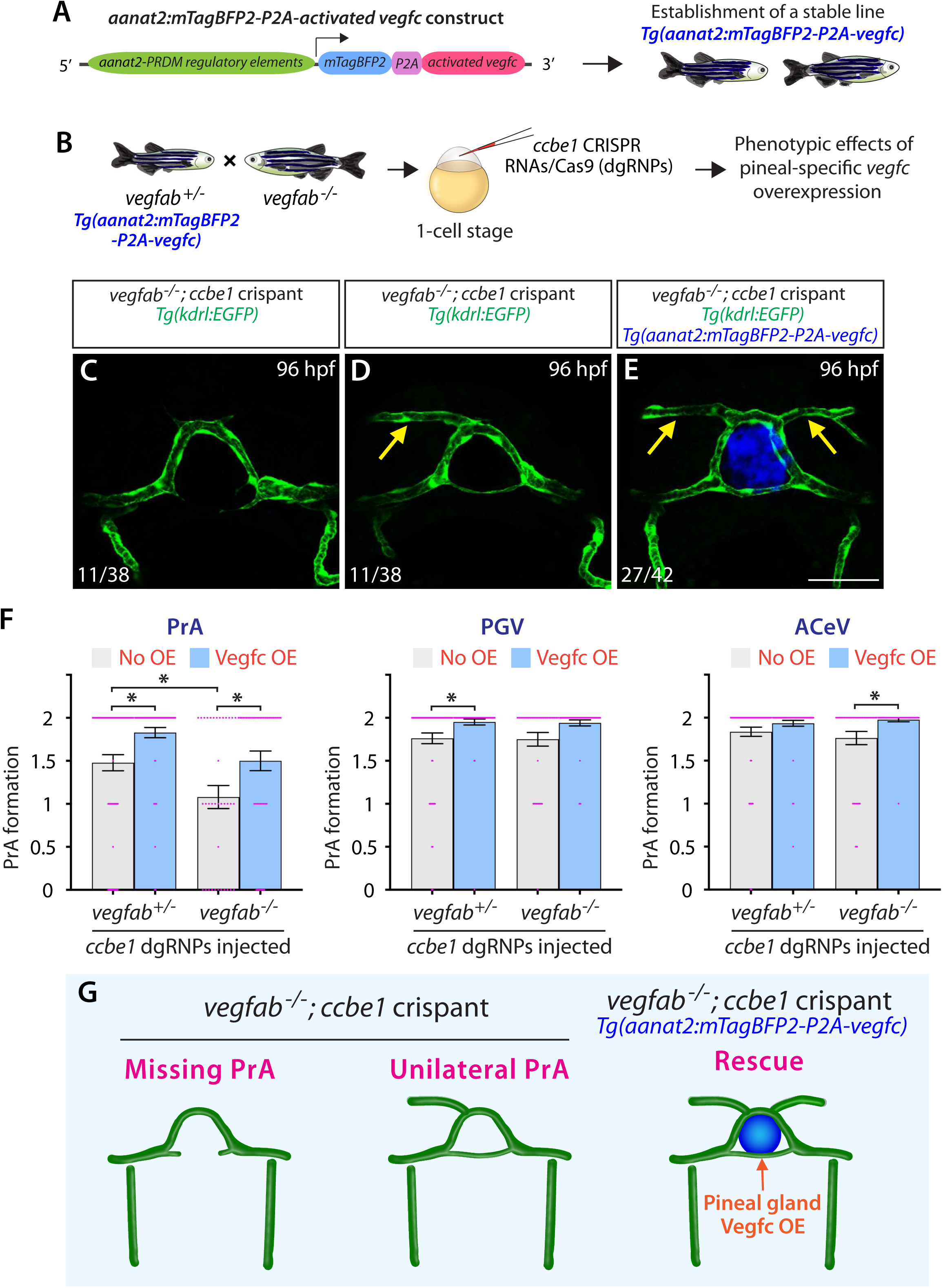
Pineal gland-specific overexpression of activated Vegfc partially rescues vascularization defects caused by combined loss of *ccbe1* and *vegfab*. (A) Schematic diagram of the donor plasmid used to generate the *Tg(aanat2:mTagBFP2-P2A-vegfc)* zebrafish line. (B) Experimental setup for the microinjection experiments shown in (**C**-**G**). (**C**-**E**) Dorsal views of the cranial vasculature at the pineal gland-dCP interface in 96 hpf *vegfab^-/-^;ccbe1* crispants (**C**-**E**) without (**C, D**) or with (**E**) the *Tg(aanat2:mTagBFP2-P2A-vegfc)* transgene. *vegfab^-/-^;ccbe1* crispants without the transgene exhibit loss of bilateral (**C**) or unilateral (**D**) PrAs, whereas the *vegfab^-/-^;ccbe1* crispant carrying the transgene exhibits bilateral PrAs (**E**). Yellow arrows indicate well-formed PrAs (**D, E**). (F) Quantification of PrA, PGV, and ACeV formation under the indicated conditions at 96 hpf. Pineal gland-specific overexpression of activated Vegfc using the *Tg(aanat2:mTagBFP2-P2A-vegfc)* transgene partially rescues vascularization defects at the pineal gland-dCP interface in *vegfab^-/-^;ccbe1* crispant larvae. Each data point represents an individual animal’s vessel formation score, and values represent mean ± SEM (\**P* < 0.05; one-way ANOVA followed by Tukey’s multiple-comparison test). (G) Schematic diagrams illustrating the vascular phenotypes at the pineal gland-dCP interface shown in (**C**-**E**). Scale bars: 50 µm in **E** for **C**-**E**.

Using these approaches, we found that pineal gland-specific overexpression of activated Vegfc significantly restored PrA formation following combined disruption of *ccbe1* and *vegfab*. While larvae deficient for both *ccbe1* and *vegfab* frequently exhibited incomplete or bilateral loss of the PrAs (Fig. 7C, D), activated Vegfc overexpression substantially increased PrA formation (Fig. 7E). Notably, the rescue was not restricted to PrA formation. Quantitative analyses further revealed significantly improved formation of the PrA, PGV, and ACeV following activated Vegfc overexpression (Fig. 7F), indicating broader restoration of the local vascular network at the pineal gland-dCP interface.

These findings demonstrate that spatially restricted expression of activated Vegfc is sufficient to partially bypass the requirement for Ccbe1 and restore vascularization at the pineal gland-dCP interface (Fig. 7G). Together with the phenotypic similarities between *ccbe1* and *vegfc* mutants, these rescue experiments provide functional evidence that Ccbe1 promotes region-specific brain vascularization through Vegfc activation. Moreover, the ability of activated Vegfc expressed specifically in the pineal gland to restore associated vascular development supports a model in which localized Vegfc activity provides a spatially restricted angiogenic signal for the formation of this cerebrovascular bed.

## Discussion

Here, we identify the Ccbe1-Adamts-Vegfc signaling axis as a crucial regulator of pineal gland and CP vascularization in the zebrafish brain. While this signaling module has been extensively studied in the context of lymphatic vessel biology (*24, 27, 32, 47*), our findings provide the first evidence that the Ccbe1-Adamts-Vegfc signaling axis is required for brain region-specific fenestrated vessel formation. The identification of this previously unrecognized role sheds light on a developmental principle underlying cerebrovascular regional heterogeneity.

Our findings propose a model in which cerebrovascular regionalization arises through the interplay between brain region-specific gene expression programs and extracellular regulation of ligand bioavailability (Fig. 8). Specifically, this work suggests that localized Ccbe1 activity spatially restricts Vegfc bioavailability during brain development. In support of this model, we showed prominent *ccbe1* expression localized to the developing pineal gland and the DMJ. Consistent with these confined expression patterns, loss of *ccbe1* and *vegfc* disrupted early vascularization of the pineal gland-dCP interface. Together with the largely preserved BBB-type vasculature in these mutants, these findings identify the Ccbe1-Vegfc axis as a core regulatory module allowing selective patterning of fenestrated cerebrovascular beds. An analogous principle has been reported in the area postrema, where locally expressed Wnt inhibitory factor-1 titrates Wnt ligand bioavailability to maintain low endothelial β-catenin signaling (*48*). Thus, multiple layers of regulation may tightly coordinate ligand bioavailability to enable cerebrovascular regionalization.

**Fig. 8.**
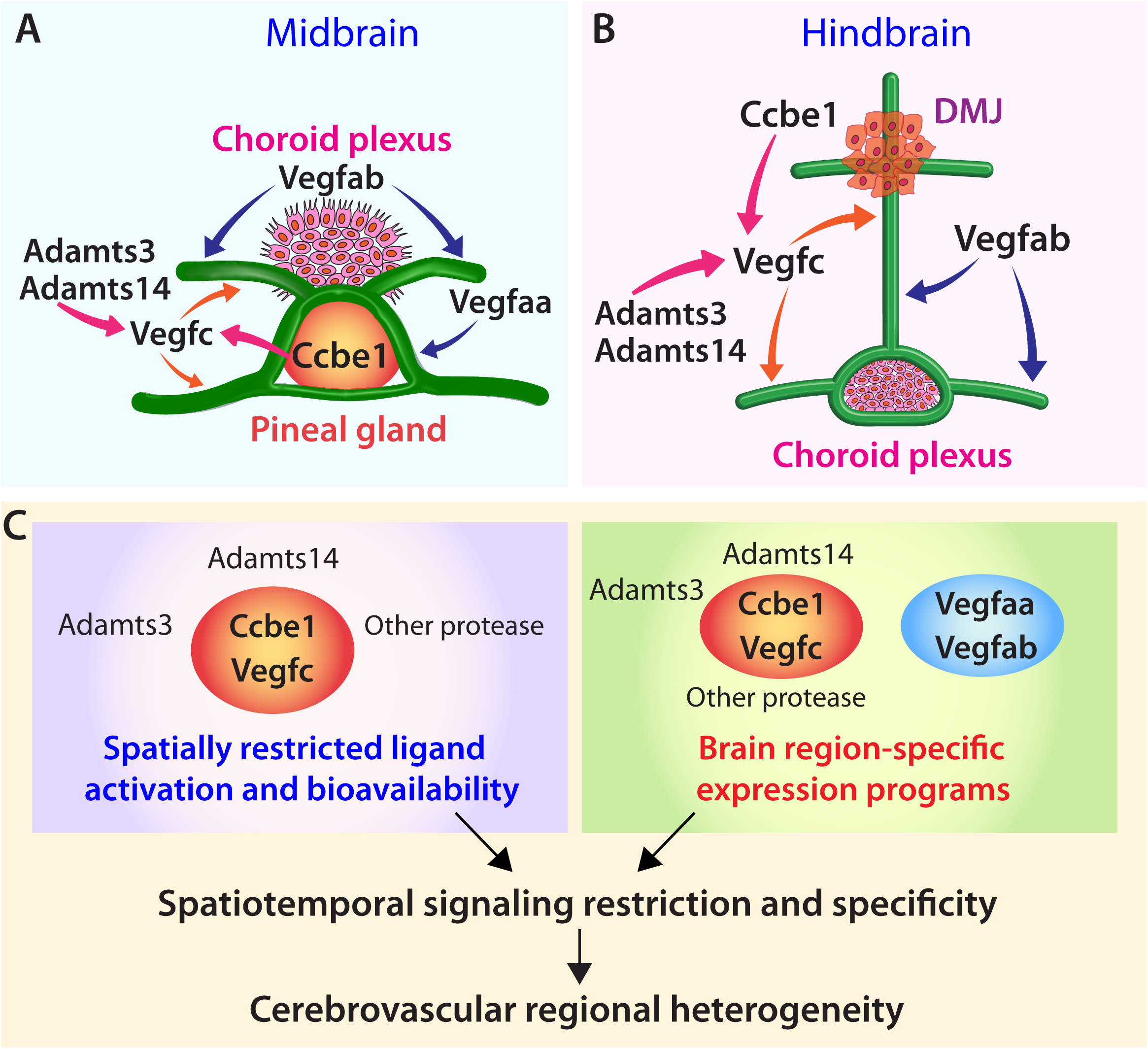
Models for spatiotemporal regulation of the Ccbe1-Adamts-Vegfc signaling axis required for region-specific fenestrated cerebrovascular development. (**a, b**) Summary diagrams illustrating the spatial organization and functional interactions of Ccbe1-Adamts-Vegfc and Vegfa signaling during vascularization at the pineal gland-dCP interface (**a**) and mCP (**b**). At the pineal gland-dCP interface, prominent *ccbe1* expression in the pineal gland provides a spatial restriction to Vegfc activation, while the distinct expression patterns of *adamts3* and *adamts14* provide an additional layer of signaling specificity. Vegfc activation cooperates with Vegfa signaling to promote local vascularization (**a**). During mCP vascularization, the Ccbe1-Vegfc signaling module cooperates with Vegfab signaling to promote formation of fenestrated vessels (**b**). Adamts3 and Adamts14 exhibit differential requirements across fenestrated vessels in this region, indicating context-dependent regulation of Vegfc activation and potential contributions of other proteases. (**c**) A conceptual model illustrating how spatially restricted ligand activation and bioavailability, along with brain region-specific expression programs, contribute to cerebrovascular regional heterogeneity. These spatially organized and temporally dynamic regulatory mechanisms provide spatiotemporal signaling specificity during cerebrovascular development.

Adamts protease-mediated activation provides an important additional layer of Vegfc signaling regulation. Previous work demonstrated that local sources of Vegfc-processing components establish signaling hubs that guide lymphatic precursor migration in the zebrafish trunk (*32*). Our findings extend this organizational principle to the cerebrovasculature, where Adamts proteases exhibit differential requirements among cerebrovascular beds. Across diverse lymphangiogenic contexts, loss of *ccbe1* closely phenocopies *vegfc* deficiency (*22, 27*), whereas the proteases responsible for Vegfc activation exhibit greater functional redundancy and context dependence (*32, 49*). Consistent with such context dependence, combined disruption of Adamts3 and Adamts14 impaired formation of the PrA and PCeV, whereas DLV formation remained largely intact. The milder cerebrovascular defects relative to Ccbe1-Vegfc signaling loss raise the possibility that additional proteases contribute to Vegfc activation. Thus, context-dependent protease activity may represent a distinct level of spatiotemporal Vegfc control that provides signaling specificity.

An unexpected observation was the remarkable capacity of the cerebrovasculature to recover. Although early vascularization of the pineal gland-dCP interface was substantially impaired by Ccbe1-Vegfc signaling loss, much of the vascular architecture was progressively restored during later development. This recovery suggests a temporally dynamic angiogenic hierarchy in which localized Vegfc signaling is particularly important during early vascular development, whereas additional angiogenic mechanisms contribute increasingly at later stages. This striking plasticity may represent developmental robustness that ensures successful formation of specialized cerebrovascular networks. Our genetic analyses identify Vegfa signaling as one such contributor, as simultaneous disruption of Vegfc and Vegfa signaling markedly enhanced pineal gland and CP vascular defects. Thus, unlike lymphatic development, which is largely dependent on the Ccbe1-Adamts-Vegfc signaling axis, fenestrated cerebrovascular development appears to involve cooperative activity of both Vegfc and Vegfa pathways. Importantly, the early onset of these cerebrovascular defects suggests that they are unlikely to arise secondarily from impaired lymphatic development, which occurs at later stages (*50–52*). Thus, distinct signaling pathways may provide complementary layers of cerebrovascular regulation.

Several important questions emerge from this study. Most notably, the developmental programs specifying localized expression of Ccbe1 and Adamts proteases within pineal gland-and CP-associated tissues remain unknown. Uncovering the mechanisms that spatially restrict this extracellular activation machinery will be an important next step toward defining how organ-specific vascular programs operate during embryogenesis. These mechanisms are particularly relevant because localized *ccbe1* expression within the meningeal mesenchymal compartment is consistent with the recent characterization of *ccbe1*-expressing fibroblasts that regulate Vegfc activation during meningeal lymphatic development (*33*). Our previous work similarly identified meningeal fibroblasts as an important cellular component regulating mCP vascularization (*13*). Controlled Ccbe1-Adamts-Vegfc activation may represent a recurring developmental strategy for organizing distinct vascular systems at brain tissue interfaces. Integration of emerging single-cell transcriptomic and chromatin accessibility datasets with genetic analyses will help identify upstream regulators responsible for these regional expression programs.

Finally, our findings may have implications beyond zebrafish cerebrovascular development. Mutations in *CCBE1* and *ADAMTS3* have been identified as causative for Hennekam syndrome (*47, 53*), a hereditary disorder characterized by lymphatic dysfunction but also associated with neurological abnormalities. Whether impaired cerebrovascular development contributes to these neurological manifestations remains unknown. Given the requirement for Ccbe1 and Vegfc during vascularization of the pineal gland and CPs, our findings raise the possibility that impaired formation of these vascular beds could influence brain development by disrupting physiological functions supported by these organs. A recent study reported that *ccbe1*-expressing meningeal fibroblasts spatially regulate Vegfc activation during the later development of meningeal lymphatic-like scavenger ECs (*33*). Together with our findings, these observations indicate that localized Ccbe1-Vegfc signaling is important at multiple stages for the development of distinct vascular systems at the brain-meningeal interface. Thus, disruption of both lymphatic and cerebrovascular development may contribute to neurological manifestations associated with human *CCBE1* deficiency. Importantly, our finding of partial PrA restoration through pineal gland-specific activated Vegfc overexpression in *ccbe1-*deficient contexts demonstrates the sufficiency of localized Vegfc activity to promote vascularization in this region.

In summary, our findings define a non-canonical role for the Ccbe1-Adamts-Vegfc axis during brain vascularization and illuminate how this signaling axis contributes to fenestrated cerebrovascular patterning. These findings support a mechanistic model in which region-specific gene expression programs and extracellular ligand bioavailability control collectively generate spatiotemporal signaling specificity within a continuous vascular network to shape cerebrovascular regional heterogeneity.

## Supporting information

Combined Supplemental Text and Figures

## Abbreviations

ACeV: Anterior cerebral vein
BBB: Blood-brain barrier
CP: Choroid plexus
CVOs: Circumventricular organs
dCP: Diencephalic choroid plexus
DLV: Dorsal longitudinal vein
DMJ: Dorsal midline junction
EC: Endothelial cell
MCeV: Middle cerebral vein
mCP: Myelencephalic choroid plexus
MsV: Mesencephalic vein
PCeV: Posterior cerebral vein
PGV: Pineal gland vessel
PrA: Prosencephalic artery
Vegfs: Vascular endothelial growth factors
Vegfrs: Vascular endothelial growth factor receptors

## Acknowledgements

We thank Dr. Yoav Gothilf for kindly providing the PRDM-aanat2 promoter-containing plasmid; Don Zeisloft and his team for zebrafish care and husbandry; Dr. Judith Drazba and her team for time-lapse and confocal imaging; and all members of the Matsuoka laboratory for discussions and comments on the manuscript. This work was supported by funding from the National Institutes of Health (R01 NS117510 and R21 NS142767) and start-up funds from the Cleveland Clinic Foundation to R.L.M.

## Author contributions

L.D.B. and R.L.M. designed the study and wrote the manuscript. L.D.B., V.T., A.G.S., F.Z.B., N.J.L., J.X.L., and A.E.L. performed loss-of-function phenotypic analyses. L.D.B., V.T., F.Z.B., and A.A.G. conducted spatial gene expression characterization. E.N.K. performed phenotypic rescue analyses. R.L.M. supervised the project. All authors have read and approved the final version of the manuscript.

## Data and materials availability

All data supporting the findings of this study are included within this manuscript. Correspondence and requests for materials should be addressed to Ryota L. Matsuoka.

## Materials and Methods

### Zebrafish husbandry and strains

All zebrafish husbandry was performed under standard conditions in accordance with institutional and national ethical and animal welfare guidelines. All zebrafish work was approved by the Cleveland Clinic Institutional Animal Care and Use Committee under protocol number 00003285. The following lines were used in this study: *Tg(kdrl:EGFP)^s843^* (*54*); *Tg(kdrl:NLS-mCherry)^is4^* (*55*); *Tg(kdrl:Has.HRAS-mcherry)^s896^* (*56*), abbreviated *Tg(kdrl:ras-mCherry)*; *Tg(kdrl:TagBFP)^mu293^* (*57*); *Tg(gata1a:DsRed)^sd2^* (*39*); *Tg(EPV.Tp1-Mmu.Hbb:Venus-Mmu.Odc1)^s940^* (*58*), abbreviated *Tg(TP1:Venus-PEST)*; *Tg(lyve1:DsRed)^nz101^* (*59*); *TgBAC(vegfaa:gal4ff)^lri96^* (*13*); *TgBAC(vegfab:gal4ff)^bns273^* (*60*); *TgBAC(vegfc:gal4ff)^bns270^* (*13*); *Tg(UAS:EGFP)^nkuasgfp1a^* (*61*); *Tg(UAS:EGFP-CAAX)^m1230^* (*62*); *vegfc^hu6410^* (*63*); *ccbe1^lri97^* (*64*); *vegfaa^bns1^* (*65*); *vegfab^bns92^* (*65*); *vegfd^bns257^* (*66*); *kdr^bns32^* (*60*); and *flt4^um131^* (*67*). Adult fish were maintained on a standard 14 h light/10 h dark daily cycle. Embryos/larvae were raised at 28.5°C. To prevent skin pigmentation, 0.003% phenylthiourea (PTU) was added beginning at 10-12 hpf for imaging. Larvae analyzed at 10 dpf were transferred to tanks containing approximately 250 mL of water supplemented with 0.003% PTU (up to 25 larvae/tank) and fed with Larval AP100 (<50 μm dry diet, Zeigler) starting at 5 dpf.

### Generation and characterization of knock-in zebrafish reporter lines

To generate knock-in zebrafish reporter lines, we used a previously established CRISPR/Cas9-based knock-in method (*68, 69*) with slight modifications. Synthetic CRISPR RNAs (crRNAs) were designed using the Custom Alt-R® CRISPR-Cas9 gRNA design tool (Integrated DNA Technologies, IDT). To generate the *ccbe1-E2A-KalTA4* knock-in line, a gRNA targeting exon 11 of *ccbe1* (gcaggaagaggaaattacct) was used. Complementary oligonucleotides (TAGGgcaggaagaggaaattacct and AAACaggtaatttcctcttcctgc) were annealed and ligated into the pT7-gRNA vector (Addgene plasmid #46759) after digestion with BsmBI (New England Biolabs, NEB). The resulting plasmid was linearized with BamHI-HF (NEB), and the gRNA was synthesized *in vitro* using the MEGAshortscript T7 Transcription Kit (Ambion) followed by purification using the RNA Clean & Concentrator-5 Kit (Zymo Research). To validate gRNA activity, approximately 1 nL of a solution containing 100 ng/μL gRNA and 2.5 μM Cas9 protein (S.p. Cas9 nuclease, V3, IDT) was injected into one-cell stage embryos. Genomic DNA isolated from injected embryos was subsequently subjected to high-resolution melt analysis (HRMA). gRNAs targeting *adamts3* (gccgacaacatcttctgagg) and *adamts14* (gcctctgactccctctggta) were designed, prepared, and validated by following the same procedures.

For generation of the *ccbe1* knock-in donor construct, a 99-bp genomic sequence containing a portion of *ccbe1* exon 11, including the gRNA recognition sequence, was used as the donor-targeting sequence, hereafter referred to as the “bait sequence”. The bait sequence was amplified by PCR using zebrafish genomic DNA as template and inserted into the *EGFPbait-E2A-KalTA4-pA* donor vector (Addgene: plasmid #61069) linearized with EcoRV using a Cold Fusion Cloning Kit (System Biosciences: MC010B-1). The following primers were used to amplify the *ccbe1* bait sequence: *ccbe1* bait sequence forward: 5’ – gcgaattctgcagatGACTTCGGATCCGGCGAAGA – 3’ *ccbe1* bait sequence reverse: 5’ – gtcctcctcctggatAACCGGCCAATCGGGATTTT – 3’

The constructed *ccbe1-E2A-KalTA4-pA* donor vector was co-injected with the *ccbe1* gRNA and Cas9 protein into one-cell stage embryos generated from incrosses of *Tg(UAS:EGFP-CAAX)^m1230I^* adults. Approximately 1 nL of an injection cocktail containing 100 ng/μL *ccbe1* gRNA, 2.5 μM Cas9 protein, 50 ng/μL *ccbe1-E2A-KalTA4-pA* donor vector, and 0.05% phenol red was injected into each embryo. Injected embryos were screened for UAS-driven EGFP-CAAX expression, and embryos exhibiting reporter expression were raised to adulthood to identify germline-transmitting founders. The established stable line was designated *KI(ccbe1-E2A-KalTA4)^lri128^*.

Donor plasmids for *adamts3* and *adamts14* knock-in generation were constructed using a similar strategy. The 108 bp and 98 bp genomic bait sequences of *adamts3* and *adamts14*, respectively, were amplified by PCR using zebrafish genomic DNA as template and inserted into the *EGFPbait-E2A-KalTA4-pA* donor vector using BamHI and EcoRV restriction enzyme sites incorporated into the forward and reverse primers, respectively. The following primers were used:

*adamts3* bait sequence forward: 5’ – attaggatccTTGTCAATCCATAGCTCGTGGAGC – 3’

*adamts3* bait sequence reverse: 5’ – attagatatcTCTCTCCACCTCAGAAGATGTTGT – 3’

*adamts14* bait sequence forward: 5’ – attaggatccGTACGGTTTCTCCAGATCTGCATC – 3’

*adamts14* bait sequence reverse: 5’ – attagatatcCGAATACTGCGGCGAATCCGTACC – 3’

The constructed *adamts3-E2A-KalTA4-pA* and *adamts14-E2A-KalTA4-pA* donor vectors were each co-injected with the corresponding *adamts3* or *adamts14* gRNA and Cas9 protein into one-cell stage embryos generated from incrosses of *Tg(UAS:EGFP-CAAX)^m1230I^* adults. Germline-transmitting founders were identified as described above for the *ccbe1* knock-in reporter line. Established stable lines were designated *KI(adamts3-E2A-KalTA4)^lri130^*and *KI(adamts14-E2A-KalTA4)^lri132^*. To characterize integration of the knock-in cassette at each targeted locus, F1 progeny from injected F0 fish were screened for KalTA4/UAS-driven EGFP-CAAX expression. Genomic DNA was extracted from reporter-positive F1 fish and subjected to PCR using a gene-specific forward primer designed within the targeted locus and a reverse primer targeting *KalTA4*. The following primers were used:

*ccbe1* exon 11 forward: 5’ – ccaccaggatcctttgacttcctg – 3’

*adamts3* exon 22 forward: 5’ – gctgctaatggatctctgaaagtc – 3’

*adamts14* exon 17 forward: 5’ – aatgagccgtgtctgggcgataag – 3’

*KalTA4* reverse: 5’-gacattcccaattgttcttcaggc – 3’

PCR products of the expected sizes were excised from agarose gels, purified, and sequenced to confirm the genomic knock-in cassette junctions and determine the precise integration sequences. Sequence analysis indicated that the resulting *ccbe1* knock-in allele is predicted to generate a Ccbe1 protein lacking the C-terminal 12 amino acids, while driving KalTA4 expression from the endogenous *ccbe1* locus. Heterozygous *ccbe1* knock-in animals exhibited no apparent vascular defects, whereas homozygous animals exhibited early vascularization defects at the pineal gland-dCP interface that were milder than those observed in stable *ccbe1^lri97^*mutant larvae, along with partial thoracic duct formation (Fig. S1). These phenotypes indicate that the knock-in allele retains partial Ccbe1 function and behaves as a hypomorphic allele. Similarly, the resulting *adamts3* and *adamts14* knock-in alleles are predicted to generate an Adamts3 protein lacking the C-terminal 7 amino acids and an Adamts14 protein lacking the C-terminal 53 amino acids, respectively. Thus, each knock-in allele enables KalTA4 expression under the control of the corresponding endogenous locus while producing a C-terminally truncated form of the encoded protein. Since individual loss of *adamts3* or *adamts14* does not produce overt cerebrovascular or trunk lymphatic defects, the extent of residual Adamts3 or Adamts14 function retained by the corresponding knock-in alleles was not rigorously assessed.

For expression analysis, while homozygous *ccbe1* knock-in animals exhibited stronger KalTA4/UAS-driven fluorescence, heterozygous animals were preferentially used when reporter intensity was sufficient, including for analyses of prominent *ccbe1* expression in the pineal gland. Homozygous animals were used selectively to increase sensitivity for detection of weaker expression domains, including meningeal and dorsal midline junction-associated expression. The overall spatial distribution of reporter expression was comparable between heterozygous and homozygous knock-in animals. Key expression patterns identified using the knock-in reporters were independently validated by RNAscope analysis of the corresponding endogenous transcripts.

### Generation of new transgenic zebrafish lines

Regulatory elements of the zebrafish *aanat2* gene, comprising the approximately 2.05 kb Pineal-Restrictive Downstream Module (PRDM) and 5′ UTR sequences of *aanat2* (*70*), were used to generate the *Tg(aanat2:mScarlet)^lri126^*and *Tg(aanat2:mTagBFP2-P2A-activated vegfc)^lri127^* transgenic lines. A transgenic construct previously used to generate the *Tg(aanat2:EGFP-2A-Δclocka-5×MYC)^tlv03^* line (*70*) was provided by Dr. Yoav Gothilf. The *gal4ff*, *mScarlet*, or *mTagBFP2-P2A-activated vegfc* transgene was inserted downstream of the *aanat2* regulatory elements by replacing the *EGFP-2A-Δclocka-5×MYC* transgene in the original construct using AgeI and NdeI restriction enzyme sites. The resulting constructs (approximately 25 pg) were injected into one-cell stage embryos together with 25 pg of transposase mRNA transcribed *in vitro* as previously described (*71*). The previously validated activated form of zebrafish Vegfc comprising amino acids 97–221 (*44*) was used to generate the *mTagBFP2-P2A-activated vegfc* construct.

### Generation and characterization of *adamts3^lri123^* and *adamts14^lri124^*mutant lines

The *adamts3^lri123^* and *adamts14^lri124^*mutant alleles were generated by CRISPR/Cas9-mediated targeted genome editing. The third exon of each gene, encoding the N-terminal prodomain region of the respective protein, was targeted. The *adamts3^lri123^* mutant allele harbors a 7 bp deletion predicted to cause a frameshift beginning at glycine residue 107, followed by 9 altered amino acids and a premature stop codon. The *adamts14^lri124^*mutant allele harbors an 8 bp deletion predicted to cause a frameshift beginning at histidine residue 69, followed by 17 altered amino acids and a premature stop codon. Both mutations are thus predicted to result in severely truncated proteins lacking the downstream metalloprotease and other functional domains. The following *adamts3* and *adamts14* crRNAs were used to generate the mutant alleles:

*adamts3* crRNA: ATTAGTGGCTCCAGGAGCCA

*adamts14* crRNA: TGCAACGGGACCAGTGATAT

Dual-guide ribonucleoprotein (dgRNP) complexes containing *adamts3* or *adamts14* crRNA, trans-activating CRISPR RNA (tracrRNA), and Cas9 protein were prepared for targeted genome editing. Injection cocktails (4 μL) containing the crRNA:tracrRNA:Cas9 RNP complexes consisted of 0.4 μL of 25 μM crRNA:tracrRNA, 0.4 μL of 25 μM Cas9 protein, 2.8 μL of H_2_O, and 0.4 μL of 0.5% phenol red solution (Sigma). Prior to microinjection, injection cocktails were incubated at 37°C for 5 min and then kept at room temperature. Approximately 1 nL of each injection cocktail was injected into the cytoplasm of one-cell stage embryos. Injected F0 fish were raised to adulthood and outcrossed to identify germline-transmitting founders, and stable mutant lines were established from F1 progeny carrying the respective deletion alleles. Genotyping of *adamts3^lri123^*and *adamts14^lri124^* mutant fish was performed by HRMA of PCR products using the following primers:

*Adamts3 lri* forward: 5’ – TTGGCAAGGAATTCCACCTCAGAC – 3’

*Adamts3 lri* reverse: 5’ – CCAGCTATTTCGATATCATCGTGC – 3’

*Adamts14 lri* forward: 5’ – CTCCCCTGGTCCAGGCTGCGTTT – 3’

*Adamts14 lri* reverse: 5’ – TGCTTGCTGCAACGGGACCAGTG – 3’

### Genotyping of mutants

Genotyping of *ccbe1^lri97^*, *vegfaa^bns1^*, and *vegfab^bns92^* mutant fish was performed by HRMA of PCR products as described previously (*13, 64*). Genotyping of *kdr^bns32^* and *flt4^um131^* mutant fish was performed by HRMA as described previously (*14*). Genotyping of *vegfc^hu6410^* and *vegfd^bns257^* mutant fish was performed by standard PCR as described previously (*13, 66*).

### High-resolution melt analysis (HRMA)

A CFX96 Touch Real-Time PCR Detection System (Bio-Rad) was used for PCR reactions and subsequent HRMA. Precision melt supermix for high-resolution melt analysis (Bio-Rad) was used for all reactions. PCR conditions were as follows: 95°C for 2 min, 46 cycles of 95°C for 10 s, and 60°C for 30 s. Following the PCR, a high-resolution melt curve was generated by collecting EvaGreen fluorescence data over a temperature range of 65–95°C. Data were analyzed using normalized derivative plots.

### Generation of F0 knockouts

dgRNPs consisting of crRNA, tracrRNA, and Cas9 protein were prepared as previously described (*64, 72*). The following *ccbe1*, *vegfab*, and *adamts3* crRNAs were used and validated for efficient disruption of their respective target loci (*64*):

*ccbe1* crRNA1: 5’ – TTCTCCTCTCGGAAAGTCCA – 3’

*ccbe1* crRNA2: 5’ – TACCCGTGCGTAAAGTCCAC – 3’

*ccbe1* crRNA3: 5’ – CTTCTGGAATACACTGACCC – 3’

*vegfab* crRNA1: 5’ – AGTGCCTACATACCCAGAGA – 3’

*vegfab* crRNA2: 5’ – TTGTTATACACCTCCATGAA – 3’

*vegfab* crRNA3: 5’ – GAAGCGTCACAATAAATAAC – 3’

*adamts3* crRNA1: 5’ – CTATGAGTAACCTAAGTGAC – 3’

*adamts3* crRNA2: 5’ – ACGTGTGAAGCGGGATGTTT – 3’

*adamts3* crRNA3: 5’ – ATTAGTGGCTCCAGGAGCCA – 3’

Injection cocktails (4 μL) containing dgRNPs targeting each gene were prepared as follows: 0.4 μL each of 25 μM crRNA1:tracrRNA, crRNA2:tracrRNA, crRNA3:tracrRNA; 1.2 μL of 25 μM Cas9 protein; 1.2 μL of H_2_O; and 0.4 μL of 0.5% phenol red solution. Injection cocktails were freshly prepared on the day of injection, and approximately 2 nL of each cocktail was injected into the cytoplasm of one-cell stage embryos.

### Immunohistochemistry

Immunohistochemistry was performed by following standard immunostaining procedures as described previously (*13*). Fish embryos/larvae were fixed in 4% paraformaldehyde (PFA) in phosphate-buffered saline (PBS), adjusted to pH 7.0, overnight at 4°C and dehydrated through immersion in methanol serial dilutions (50%, 75%, and 100%, followed by two additional washes in 100%, 10 min each) at room temperature (RT). Dehydrated samples were stored in 100% methanol at - 20°C until use. Samples were rehydrated through immersion in methanol serial dilutions (50% and 25%, 10 min each) at RT and washed briefly three times in 1% PBST (1% Triton X-100 in 0.1M PBS) Samples were then permeabilized with 10 µg/ml Proteinase K in 1% PBST for 15 min at RT and blocked for 2-4 h at RT in blocking solution containing 0.5% bovine serum albumin in 1% PBST.

The following primary antibodies were used: chicken anti-GFP (GFP-1010, Aves Labs, 1:1000) and rabbit anti-DsRed (#632496, Clontech, 1:300). Following overnight incubation with primary antibodies at 4°C, samples were washed, incubated with appropriate secondary antibodies (1:500) overnight at 4°C, washed again, and processed for imaging as described previously (*13*).

### RNAscope *in situ* hybridization

RNAscope assays were performed on *Tg(kdrl:EGFP)^s843^*embryos collected at 48 hpf by following procedures described previously (*14*). Samples were fixed, dehydrated, and rehydrated as described above for immunohistochemistry. Permeabilization was performed using Protease Plus (#322331; Advanced Cell Diagnostics, ACD) at 40°C for 3 min in the HybEZ^TM^ II hybridization system (#321711, ACD). After permeabilization, samples were washed three times in 1% PBST at RT. Gene-specific probes were designed and synthesized by ACD and prepared according to the manufacturer’s instructions. Probes specific for *ccbe1* (#815251-C3, 50X), *adamts3* (#416611-C2, 50X), and *adamts14* (#1205121-C2, 50X) were freshly diluted to 1X in probe diluent (#300041, ACD) for each assay. Probe hybridization was performed at 40°C for 2 h. During hybridization, 1X wash buffer was prepared from 50X stock (#310091; ACD) using H₂O and pre-warmed to 40°C.

Following hybridization, signal detection was performed using the RNAscope Multiplex Fluorescent Detection Kit v2 (#323110, ACD). Samples were washed three times in pre-warmed wash buffer at 40°C for 10 min each. Pre-warmed amplification reagents Amp 1, Amp 2, Amp 3 were then applied sequentially at 40°C for 30, 30, and 15 min, respectively, with two intervening washes in wash buffer at RT for 5 min each. Following Amp3 incubation, samples were washed twice in wash buffer at RT for 5 min each and kept overnight at RT in 5X SSC (750 mM NaCl, 75 mM sodium citrate, pH 7.0).

The following day, samples were washed once in wash buffer for 5 min at RT and incubated with the appropriate pre-warmed HRP-conjugated reagent (HRP-C1, HRP-C2, or HRP-C3) at 40°C for 15 min, followed by two washes in wash buffer at RT for 5 min each. Fluorescent signal was developed using TSA Vivid Fluorophore 650 (#323273, ACD) diluted at 1:750 in Multiplex TSA buffer (#322810, ACD) at 40°C for 30 min. Samples were subsequently protected from prolonged light exposure to preserve fluorescence. After two washes in wash buffer at RT for 5 min each, samples were incubated with pre-warmed Multiplex FL v2 HRP Blocker (#323107, ACD) at 40°C for 15 min. Following HRP blocking, samples were washed once in 1% PBST at RT and incubated with the anti-GFP antibody described above for at least 4 h at RT. After three washes in 1% PBST at RT for 20 min each, samples were incubated with the appropriate secondary antibody overnight at 4 °C. After six washes in 1% PBST at RT for 15 min each, samples were processed for imaging.

### Confocal and stereo microscopy

Fluorescence imaging of live and fixed zebrafish samples was conducted using a Leica TCS SP8 confocal laser scanning microscope. Fish embryos and larvae were anesthetized with a low dose of tricaine, embedded in 1% low melt agarose in glass-bottom Petri dishes (MatTek), and imaged using a 25X water-immersion objective lens. Image acquisition and analysis were performed using LAS X software (Version 3.7.0.20979). Brightfield images of anesthetized fish were captured using a Nikon SMZ-18 stereomicroscope. Image acquisition and analysis for brightfield imaging were conducted using NIS-Elements BR Imaging software (Version 5.10.01).

### Time-lapse cell tracking of erythrocytes and brain hemorrhage analysis

*Tg(kdrl:EGFP)^s843^;Tg(gata1a:DsRed)^sd2^* larvae were used for live imaging of blood flow and analysis of local brain hemorrhage at 96 hpf. For blood-flow analysis, DsRed-positive erythrocytes were imaged using a Leica TCS SP8 confocal microscope equipped with a high-speed resonant scanner. Images were acquired at approximately 32 ms intervals within a single *z*-plane, enabling individual erythrocytes to be tracked through the local cerebrovascular network. A maximum-intensity projection of a *z*-stack image of the corresponding cerebrovascular network visualized by the *Tg(kdrl:EGFP)* reporter was acquired once and overlaid onto sequential single-*z*-plane time-lapse images of DsRed-positive erythrocytes using Fiji software. The direction of blood flow was determined by tracking the movement of DsRed-positive erythrocytes through the vessels of interest at the pineal gland-dCP interface and mCP.

For local brain hemorrhage analysis, *Tg(kdrl:EGFP);Tg(gata1a:DsRed)* larvae were examined at 96 hpf for abnormal accumulation of DsRed-positive erythrocytes outside the vascular lumen. Hemorrhage was assessed in defined regions surrounding the pineal gland-dCP interface and mCP, and the frequency of larvae exhibiting local hemorrhage was determined for each genotype. Vascular architecture was simultaneously visualized using the *Tg(kdrl:EGFP)* reporter to relate extravascular erythrocyte accumulation to local vascularization defects.

### Quantification of PrA, PGV, ACeV, DLV, PCeV, MsV, and MCeV formation

Quantification was performed using fish carrying the *Tg(kdrl:EGFP)^s843^* reporter. Fish larvae at the indicated developmental stages were analyzed for the presence or absence of the DLV. To assess bilateral formation of the PrA, PGV, ACeV, PCeV, MsV, and MCeV, the following scoring criteria were applied: Score 2, both bilateral vessels were fully formed; Score 1.5, one vessel was fully formed while the other was partially formed; Score 1, both vessels were partially formed or one vessel was fully formed while the other was absent; Score 0.5, one vessel was partially formed while the other was absent; and Score 0, both bilateral vessels were absent.

For early embryos examined at 32 and 48 hpf, formation of the vascular unit composed of the bilateral PrA, PGV, and ACeV was classified as either fully formed or incompletely formed. A vascular unit was considered fully formed when the PrA, PGV, and ACeV were present bilaterally and established the expected continuous vascular connections. Any vascular unit exhibiting partial formation or absence of one or more of these vessels was classified as incompletely formed.

### Quantification of pineal gland size

Pineal gland size was quantified in fish carrying the *Tg(EPV.Tp1-Mmu.Hbb:Venus-Mmu.Odc1)^s940^* line at 32 hpf. The anatomical location and boundaries of the pineal gland were identified based on the spatial distribution of Notch reporter-positive cells in combination with differential interference contrast (DIC) imaging. Specifically, maximum-intensity projection images of the Notch reporter signal covering the pineal gland along its entire dorsoventral axis were used together with selected *z*-plane DIC images that clearly visualized the anatomical boundaries of the pineal gland. A region of interest (ROI) was manually delineated along the outer boundary of the pineal gland using Fiji software, and the enclosed area was measured to determine pineal gland size. The same criteria were applied to all samples across experimental groups.

### Quantification of thoracic duct (TD) formation

Fish carrying *Tg(lyve1:DsRed)^nz101^* were used for quantification of TD formation. TD formation was scored from lateral-view confocal images spanning three complete somites. Within each somite, TD formation was assigned a score of 1 when the TD fully connected the two adjacent intersegmental vessels (ISVs) and a score of 0 when this connection was incomplete or absent. Because each image encompassed three somites, the scores were summed to generate a total score ranging from 0 to 3 for each fish.

## Statistical analysis

Statistical differences in mean values among multiple groups were determined using a one-way analysis of variance (ANOVA) followed by Tukey’s multiple comparison test. Statistical differences in mean values between two groups were determined using a two-tailed Student’s *t*-test. Fisher’s exact test was applied to assess statistical differences in the penetrance of observed phenotypes. Statistical analyses were conducted using GraphPad Prism version 11.0.2. The criterion for statistical significance was set at *P* < 0.05. Error bars represent SEM.

