## Supplementary material for "The lymphangiogenic Ccbe1-Adamts-Vegfc axis shapes cerebrovascular regionalization via selective patterning of fenestrated vascular beds": Combined Supplemental Text and Figures

#### Supplemental Figure Legends

**Fig. S1. Characterization of *ccbe1* knock-in heterozygous and homozygous lines.**
(**A**-**C**) Dorsal views of 32 hpf *ccbe1* knock-in heterozygous (**A**) and homozygous (**B**, **C**) embryos carrying the *Tg(UAS:EGFP);Tg(kdrl:ras-mCherry)* reporters, showing EGFP expression domains in the pineal gland and associated blood vessels in this region. While the *ccbe1* knock-in heterozygous embryo displayed well-formed vasculature in this brain region (**A**), homozygous embryos (**B**, **C**) exhibited variable vascular phenotypes, with either incomplete (**B**) or well-formed (**C**) vascularization. Impaired vascular formation at the pineal gland-dCP interface is indicated by cyan asterisks (**B**).
(**D**) Quantification of vascular formation at the pineal gland-dCP interface in 32 hpf WT, *ccbe1* knock-in heterozygous and homozygous, and *ccbe1* homozygous mutant embryos (numbers of animals examined per genotype are listed in the panel). WT and *ccbe1* homozygous mutant data are re-presented from **Fig. 2K** for direct comparison with the knock-in genotypes. While *ccbe1* knock-in heterozygous embryos exhibited vascular phenotypes at this interface similar to WT animals, homozygous knock-in embryos displayed impaired vascular formation that was significantly milder than that observed in *ccbe1* homozygous mutants, indicating that the *ccbe1* knock-in allele behaves as a hypomorph. $^{**}P < 0.01$ and $^{****}P < 0.0001$; Fisher's exact test. NS: not significant.
(**E**-**G**) Lateral views of 6 dpf *ccbe1* knock-in heterozygous (**E**) and homozygous (**F**, **G**) larvae carrying the *Tg(UAS:EGFP-CAAX);Tg(lyve1:DsRed)* reporters, showing venous and lymphatic vessels along with EGFP-CAAX expression domains in the ventral trunk. While the *ccbe1* knock-in heterozygous larva displayed a well-formed thoracic duct (TD) dorsal to the posterior cardinal vein (PCV) (**E**, arrows), homozygous larvae (**F**, **G**) exhibited variable vascular phenotypes, with either nearly absent TD formation (**F**) or a well-formed TD (**G**). Impaired TD formation is indicated by cyan asterisks (**F**).
(**H**) Quantification of TD formation in the ventral trunk of 6 dpf WT, *ccbe1* knock-in heterozygous and homozygous larvae. While *ccbe1* knock-in heterozygous larvae exhibited TD formation similar to WT animals, homozygous knock-in larvae displayed significantly impaired TD formation. Given that *ccbe1* homozygous mutants have been reported to lack TD formation, the impaired TD formation observed in *ccbe1* homozygous knock-in larvae was substantially milder, further supporting the hypomorphic nature of the *ccbe1* knock-in allele. Each data point represents an individual animal's TD formation score, and values represent mean ± SEM ($^{***}P < 0.001$; one-way ANOVA followed by Tukey's multiple-comparison test; NS: not significant).
Scale bars: 50 µm in **C** for **A**-**C**; **G** for **E**-**G**.

**Fig. S2. Ccbe1 cooperates with Vegfa and Vegfd to promote region-specific fenestrated vessel formation at the pineal gland-choroid plexus interface and in the hindbrain mCP.**
(**A**) Schematic dorsal view of the vascular anatomy at the pineal gland-dCP interface at 10 dpf.
(**B**-**F**) Dorsal views of the cranial vasculature at the pineal gland-dCP interface in 10 dpf WT (**B**), *ccbe1*$^{-/-}$ (**C**), *vegfab*$^{-/-}$ (**D**), *ccbe1*$^{-/-}$*;vegfab*$^{-/-}$ (**E**), and *ccbe1*$^{-/-}$*;vegfab*$^{-/-}$*;vegfd*$^{-/-}$ (**F**) larvae visualized by the *Tg(kdrl:*EGFP*)* transgene. Yellow arrows point to the PrA, blue arrows to the PGV, and orange arrows to the ACeV.
(**G**) Quantification of PrA, PGV, and ACeV formation in the indicated genotypes at 10 dpf (numbers of animals examined per genotype are listed in the panel). PrA formation defects resulting from the genetic interaction between *ccbe1* and *vegfab* persist through 10 dpf. Additional loss of *vegfd* preferentially enhances impaired PGV formation, supporting a region- and vessel-specific contribution of Vegfd signaling.
(**H**) Schematic dorsal view of the vascular anatomy near the mCP at 10 dpf, indicating the blood vessels examined in this region.
(**I**-**M**) Dorsal views of the cranial vasculature near the mCP in 10 dpf WT (**I**), *ccbe1*$^{-/-}$ (**J**), *vegfab*$^{-/-}$ (**K**), *ccbe1*$^{-/-}$*;vegfab*$^{-/-}$ (**L**), and *ccbe1*$^{-/-}$*;vegfab*$^{-/-}$*;vegfd*$^{-/-}$ (**M**) larvae visualized by the *Tg(kdrl:*EGFP*)* transgene,

showing DLV loss in *vegfab*$^{-/-}$ larva and complete loss of the DLV and bilateral PCeVs in *ccbe1*$^{-/-}$*;vegfab*$^{-/-}$ and *ccbe1*$^{-/-}$*;vegfab*$^{-/-}$*;vegfd*$^{-/-}$ larvae. Yellow arrows point to the DLV and white arrows to the PCeV.
(**N**) Percentage of 10 dpf larvae of the indicated genotypes with or without the DLV (numbers of animals examined per genotype are listed in the panel). Combined loss of *ccbe1* and *vegfab* causes a strong, persistent DLV formation defect that does not recover by 10 dpf. Additional loss of *vegfd* further enhances this phenotype, with no triple-mutant larvae forming the DLV. $*P < 0.05$, $***P < 0.001$, and $****P < 0.0001$; Fisher's exact test. NS: not significant.
(**O**) Quantification of PCeV, MsV, and MCeV formation in the indicated genotypes at 10 dpf. Combined loss of *ccbe1* and *vegfab* strongly impairs PCeV formation, whereas MsV and MCeV formation remains largely unaffected. Additional loss of *vegfd* further enhances PCeV defects, with no triple-mutant larvae forming bilateral PCeVs. In panels (**G** and **O**), each data point represents an individual animal's vessel formation score, and values represent mean ± SEM ($**P < 0.01$, $***P < 0.001$, and $****P < 0.0001$; one-way ANOVA followed by Tukey's multiple-comparison test; NS: not significant).
Scale bars: 50 μm in **F** for **B**-**F**; **M** for **I**-**M**.

**Fig. S3. Combined loss of Ccbe1 and Vegfc causes only limited enhancement of region-specific fenestrated vessel formation defects.**
(**A**) Experimental setup for the cranial vascular phenotypic analysis shown in (**B**-**F**).
(**B**, **C**) Dorsal views of the cranial vasculature at the pineal gland-dCP interface in 96 hpf WT (**B**) and *ccbe1*$^{-/-}$*;vegfc*$^{-/-}$ (**C**) larvae visualized by the *Tg(kdrl:*EGFP*)* transgene, showing no enhanced vascular defects in this region in *ccbe1*$^{-/-}$*;vegfc*$^{-/-}$ larvae.
(**D**, **E**) Dorsal views of the cranial vasculature near the mCP in 96 hpf WT (**D**) and *ccbe1*$^{-/-}$*;vegfc*$^{-/-}$ (**E**) larvae visualized by the *Tg(kdrl:*EGFP*)* transgene, showing impaired PCeV formation (cyan asterisks) in *ccbe1*$^{-/-}$*;vegfc*$^{-/-}$ larvae.
(**F**) Quantification of PrA, PGV, and ACeV formation in the indicated genotypes at 96 hpf (numbers of animals examined per genotype are listed in the panel). Combined loss of *ccbe1* and *vegfc* does not significantly enhance vascular formation defects in this region.
(**G**) Percentage of 96 hpf larvae of the indicated genotypes with or without the DLV. Combined loss of *ccbe1* and *vegfc* does not significantly enhance DLV formation defects. Statistical significance was assessed by Fisher's exact test (NS: not significant).
(**H**) Quantification of PCeV, MsV, and MCeV formation in the indicated genotypes at 96 hpf. Combined loss of *ccbe1* and *vegfc* mildly enhances PCeV formation defects, whereas MsV and MCeV formation remains unaffected. In panels (**F** and **H**), each data point represents an individual animal's vessel formation score, and values represent mean ± SEM ($*P < 0.05$, $***P < 0.001$, and $****P < 0.0001$; one-way ANOVA followed by Tukey's multiple-comparison test; NS: not significant).
Scale bars: 50 μm in **B** for **B**, **C**; **D** for **D**, **E**.

**Fig. S4. Ultrastructural analysis of 10 dpf WT and DLV-lacking *vegfab*$^{-/-}$ larvae reveals disrupted mCP epithelial organization in mutants.**
(**A**, **A'**) Dorsal view of a 10 dpf WT head section stained with toluidine blue. Magnified view of the boxed area in (**A**) indicates the location of the mCP (**A'**, dashed lines).
(**B**) TEM image of the corresponding mCP and surrounding tissue at 10 dpf. The mCP exhibits morphological characteristics of epithelial cell populations (light gray) compared with the darker surrounding tissues.
(**C**-**H**) TEM images of the mCP and surrounding tissues in 10 dpf WT (**C**, **E**, **G**) and DLV-lacking *vegfab*$^{-/-}$ (**D**, **F**, **H**) larvae. WT larvae exhibit a well-organized epithelial cell layer characterized by centrally located cell bodies and microvilli extending toward the surrounding tissues. In contrast, the *vegfab*$^{-/-}$ larva lacking the DLV exhibits disrupted epithelial cell organization, including apparently

misoriented microvilli (cyan arrows in **F**, **H**) and ectopic swollen-like cellular structures (orange arrows in **F**, **H**). CPE: choroid plexus epithelial cell; MV: microvilli.
Scale bars: 5 μm in **B**; 2 μm in **C** for **C**, **D**; 2 μm in **E** for **E**, **F**; 1 μm in **G** for **G**, **H**.

**Fig. S5. Knock-in strategies and validation of *KI(adamts3-E2A-KalTA4)*- and *KI(adamts14-E2A-KalTA4)*-driven reporter expression by comparison with endogenous mRNA expression.**
(**A**) Strategy for generating an *adamts3* knock-in zebrafish line and genomic characterization of the established line.
(**B**) Strategy for generating an *adamts14* knock-in zebrafish line and genomic characterization of the established line.
(**C**, **D**) Side-by-side comparison of a 48 hpf heterozygous *adamts3* knock-in embryo carrying the *Tg(UAS:EGFP-CAAX);Tg(kdrl:ras-mCherry)* reporters (**C**) and *adamts3* mRNA expression (**D**), showing consistent expression patterns in the developing head.
(**E**, **F**) Side-by-side comparison of a 48 hpf heterozygous *adamts14* knock-in embryo carrying the *Tg(UAS:EGFP-CAAX);Tg(kdrl:ras-mCherry)* reporters (**E**) and *adamts14* mRNA expression (**F**), showing consistent expression patterns in the developing head, most prominently in the otic vesicle.
Scale bars: 50 μm in **F** for **C**-**F**.

**Fig. S6. Side-by-side comparison of *ccbe1*, *adamts3*, and *adamts14* knock-in reporter expression in 55 and 96 hpf embryos and larvae.**
(**A**-**C**) Lateral views of 55 hpf *ccbe1* (**A**), *adamts3* (**B**), and *adamts14* (**C**) knock-in homozygous embryonic heads carrying the *Tg(UAS:EGFP-CAAX);Tg(kdrl:ras-mCherry)* reporters, showing the spatial distributions of reporter expression in the developing brain.
(**D**-**F**) Lateral views of 96 hpf *ccbe1* (**D**), *adamts3* (**E**), and *adamts14* (**F**) knock-in homozygous larvae carrying the *Tg(UAS:EGFP-CAAX);Tg(kdrl:ras-mCherry)* reporters, showing reporter expression throughout the body.
Scale bars: 50 μm in **C** for **A**-**C**; 500 μm in **D** for **D**-**F**.

**Fig. S7. Generation and validation of *adamts3$^{lri123}$* and *adamts14$^{lri124}$* mutant alleles.**
(**A**) Predicted protein structures of zebrafish WT Adamts3 and Adamts3$^{lri123}$ (left), and WT Adamts14 and Adamts14$^{lri124}$ (right). Both WT Adamts proteins contain a signal peptide (SP), prodomain, catalytic domain, disintegrin-like domain, thrombospondin type I repeat (TSR), cysteine-rich domain, spacer domain, three additional TSRs, and a procollagen N-propeptidase domain from the N- to C-terminus. Adamts3$^{lri123}$ and Adamts14$^{lri124}$ are predicted to retain only the SP and a portion of the prodomain, followed by short missense amino-acid stretches generated by frameshift mutations before premature stop codons. Arrows indicate the regions within the prodomain targeted by genome editing, and red lines indicate the predicted missense amino-acid sequences downstream of the mutations.
(**B**, **C**) Sequence alignment of the targeted *adamts3* region from the *adamts3$^{+/+}$* and *adamts3$^{lri123/lri123}$* alleles shows a CRISPR-induced 7-bp deletion (GGAGCCA; c.319_325del) (**B**, left), which was confirmed by Sanger sequencing chromatograms of PCR products generated from *adamts3$^{+/+}$* and *adamts3$^{lri123/lri123}$* genomic DNA (**C**, left). Similarly, analysis of the targeted *adamts14* region identified an 8-bp deletion (CACCGATA; c.205_212del) in the *adamts14$^{lri124/lri124}$* allele (**B**, right), which was confirmed by Sanger sequencing chromatograms (**C**, right).
(**D**) High-resolution melt analysis (HRMA) used to distinguish WT, heterozygous, and homozygous *adamts3$^{lri123}$* (left) and *adamts14$^{lri124}$* (right) genotypes. Melting curves for three independent fish of each genotype are shown in the indicated colors.

**Fig. S8. Individual loss of *adamts3* or *adamts14* does not cause significant brain vascularization defects in either WT or *vegfab* mutant backgrounds.**

(**A**-**C**) Quantification of cerebrovascular formation at 10 dpf in WT, *adamts3*$^{-/-}$, *adamts14*$^{-/-}$, *vegfab*$^{-/-}$, *adamts3*$^{-/-}$;*vegfab*$^{-/-}$, and *adamts14*$^{-/-}$;*vegfab*$^{-/-}$ larvae. Numbers of animals examined for each genotype are indicated in the figure.
(**A**) Quantification of PrA, PGV, and ACeV formation. No significant differences were observed among the indicated genotypes.
(**B**) Percentage of 10 dpf larvae with or without the DLV. Individual loss of *adamts3* or *adamts14* did not cause DLV formation defects. Furthermore, combined loss of *vegfab* and either *adamts3* or *adamts14* did not significantly enhance the DLV formation defects observed in *vegfab* single mutants. Statistical significance was assessed by Fisher's exact test (NS: not significant).
(**C**) Quantification of PCeV, MsV, and MCeV formation. No significant differences were observed among the indicated genotypes. NS: not significant.
In panels (**A** and **C**), each data point represents an individual animal's vessel formation score, and values represent mean ± SEM (NS: not significant; one-way ANOVA followed by Tukey's multiple-comparison test).

**Movie S1. Time-lapse tracking of red blood cells at the pineal gland-dCP interface in a 96 hpf WT larva.**
Time-lapse tracking of red blood cell movement in a 96 hpf WT larva visualized by the *Tg(kdrl:*EGFP*)* and *Tg(gata1a:DsRed)* reporters, showing directional red blood cell movement from the PrA to the ACeV through the PGV.

**Movie S2. Time-lapse tracking of red blood cells at the pineal gland-dCP interface in a 96 hpf *ccbe1*$^{-/-}$;*vegfab*$^{-/-}$ larva.**
Time-lapse tracking of red blood cell movement in a 96 hpf *ccbe1*$^{-/-}$;*vegfab*$^{-/-}$ larva visualized by the *Tg(kdrl:*EGFP*)* and *Tg(gata1a:DsRed)* reporters, showing abnormal blood flow from one side of the ACeV to the other in the absence of bilateral PrAs.

### Figure S1

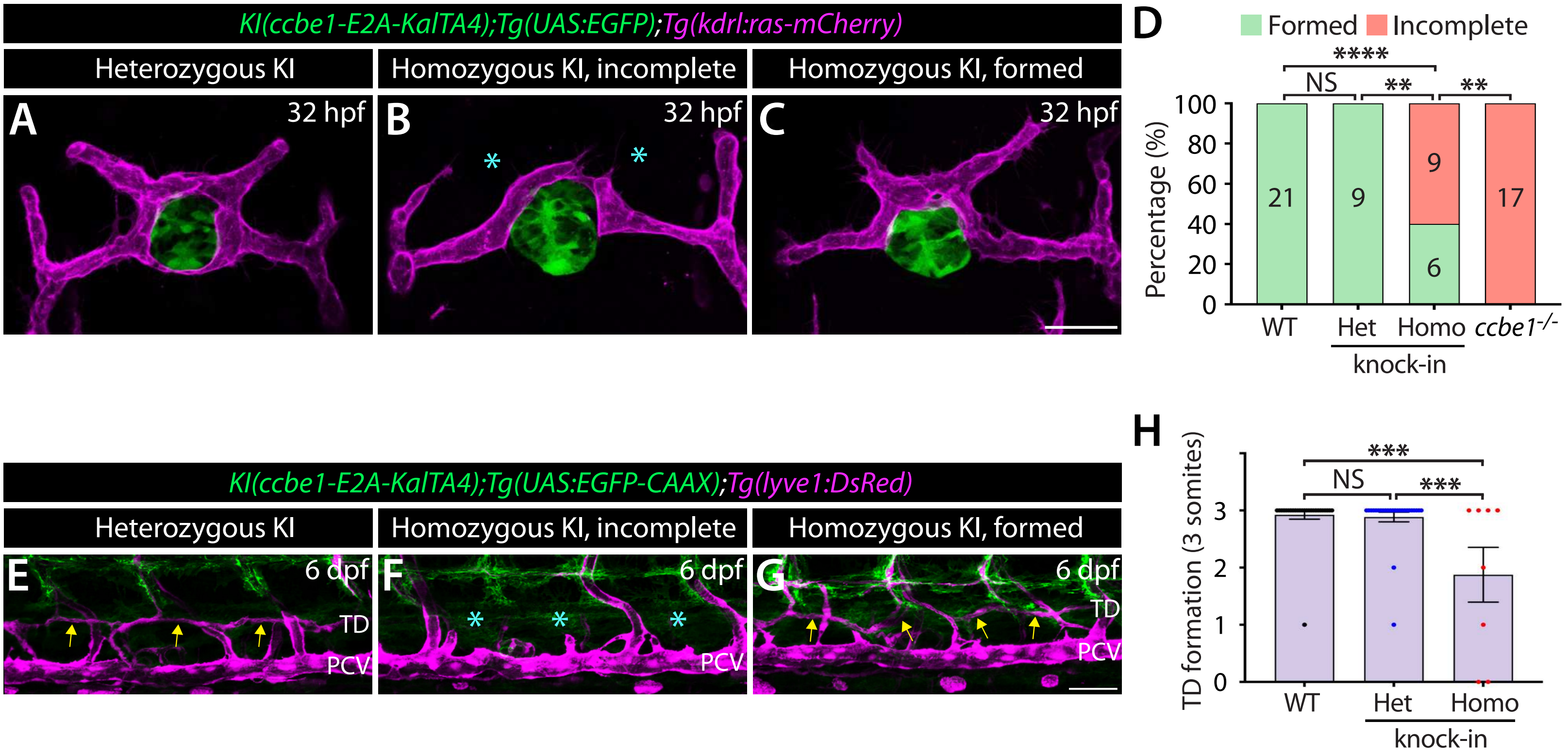

KI(ccbe1-E2A-KalTA4);Tg(UAS:EGFP);Tg(kdrl:ras-mCherry)
Heterozygous KI
Homozygous KI, incomplete
Homozygous KI, formed
A
B
C
32 hpf
D
Formed
Incomplete
Percentage (%)
100
80
60
40
20
0
21
9
9
6
17
NS
****
**
WT
Het
Homo
ccbe1-/-
knock-in
KI(ccbe1-E2A-KalTA4);Tg(UAS:EGFP-CAAX);Tg(lyve1:DsRed)
E
F
G
6 dpf
TD
PCV
H
TD formation (3 somites)
***
3
2
1

Figure S2

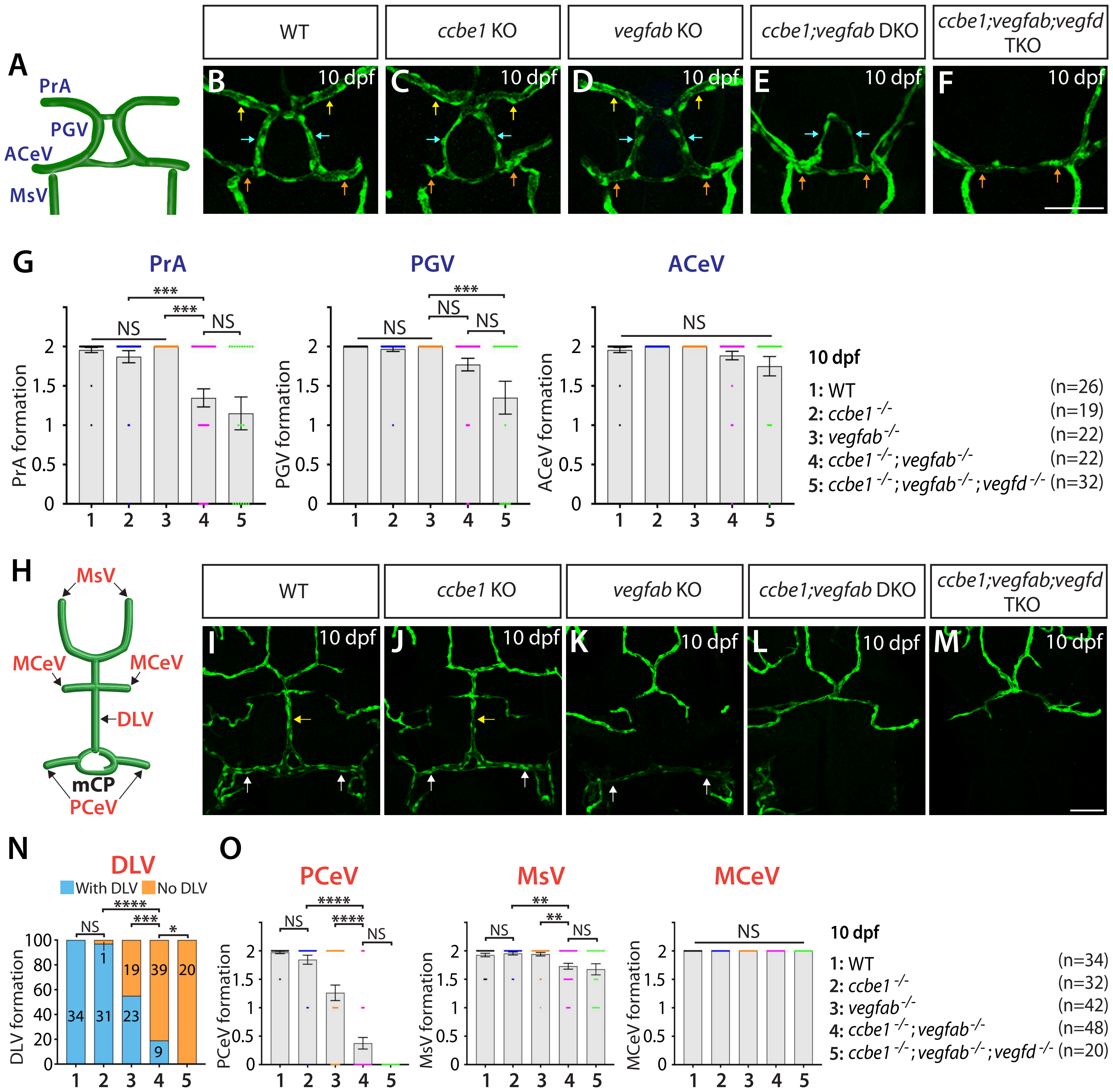
A
PrA
PGV
ACeV
MsV
WT
ccbe1 KO
vegfab KO
ccbe1;vegfab DKO
ccbe1;vegfab;vegfd TKO
B 10 dpf
C 10 dpf
D 10 dpf
E 10 dpf
F 10 dpf
G
PrA
PGV
ACeV
PrA formation
PGV formation
ACeV formation
NS
***
10 dpf
1: WT (n=26)
2: ccbe1 -/- (n=19)
3: vegfab -/- (n=22)
4: ccbe1 -/- ; vegfab -/- (n=22)
5: ccbe1 -/- ; vegfab -/- ; vegfd -/- (n=32)
H
MsV
MCeV
DLV
mCP
PCeV
I 10 dpf
J 10 dpf
K 10 dpf
L 10 dpf
M 10 dpf
N
DLV
With DLV
No DLV
DLV formation
O
PCeV
MsV
MCeV
PCeV formation
MsV formation
MCeV formation
NS
*
**
***
****
10 dpf
1: WT (n=34)
2: ccbe1 -/- (n=32)
3: vegfab -/- (n=42)
4: ccbe1 -/- ; vegfab -/- (n=48)
5: ccbe1 -/- ; vegfab -/- ; vegfd -/- (n=20)

### Figure S3

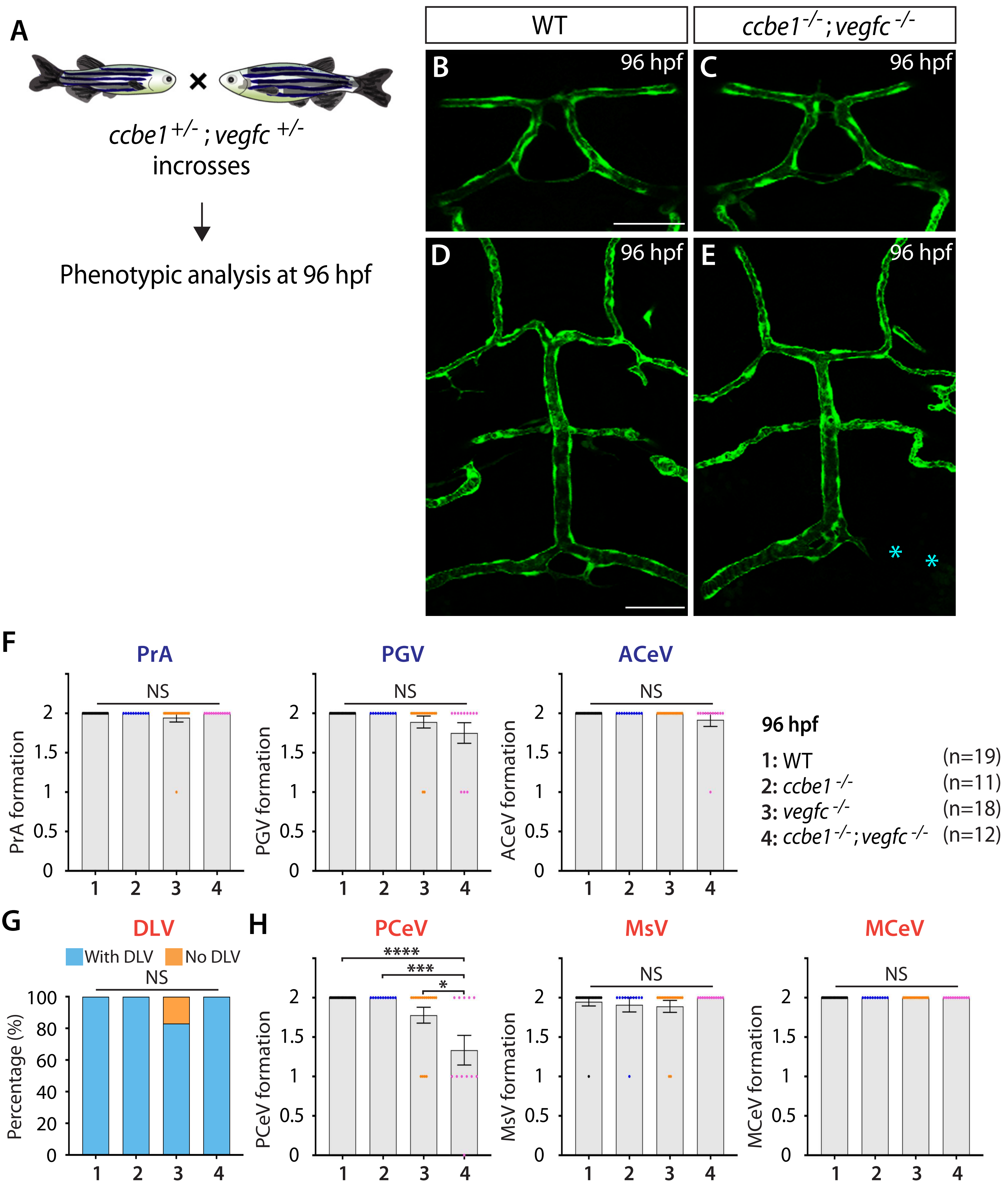
A
ccbe1 +/- ; vegfc +/-
incrosses
Phenotypic analysis at 96 hpf
WT
ccbe1 -/- ; vegfc -/-
B 96 hpf
C 96 hpf
D 96 hpf
E 96 hpf
F
PrA
PGV
ACeV
NS
PrA formation
PGV formation
ACeV formation
96 hpf
1: WT (n=19)
2: ccbe1 -/- (n=11)
3: vegfc -/- (n=18)
4: ccbe1 -/- ; vegfc -/- (n=12)
G
DLV
With DLV
No DLV
Percentage (%)
H
PCeV
MsV
MCeV
****
***
*
PCeV formation
MsV formation
MCeV formation

Figure S4

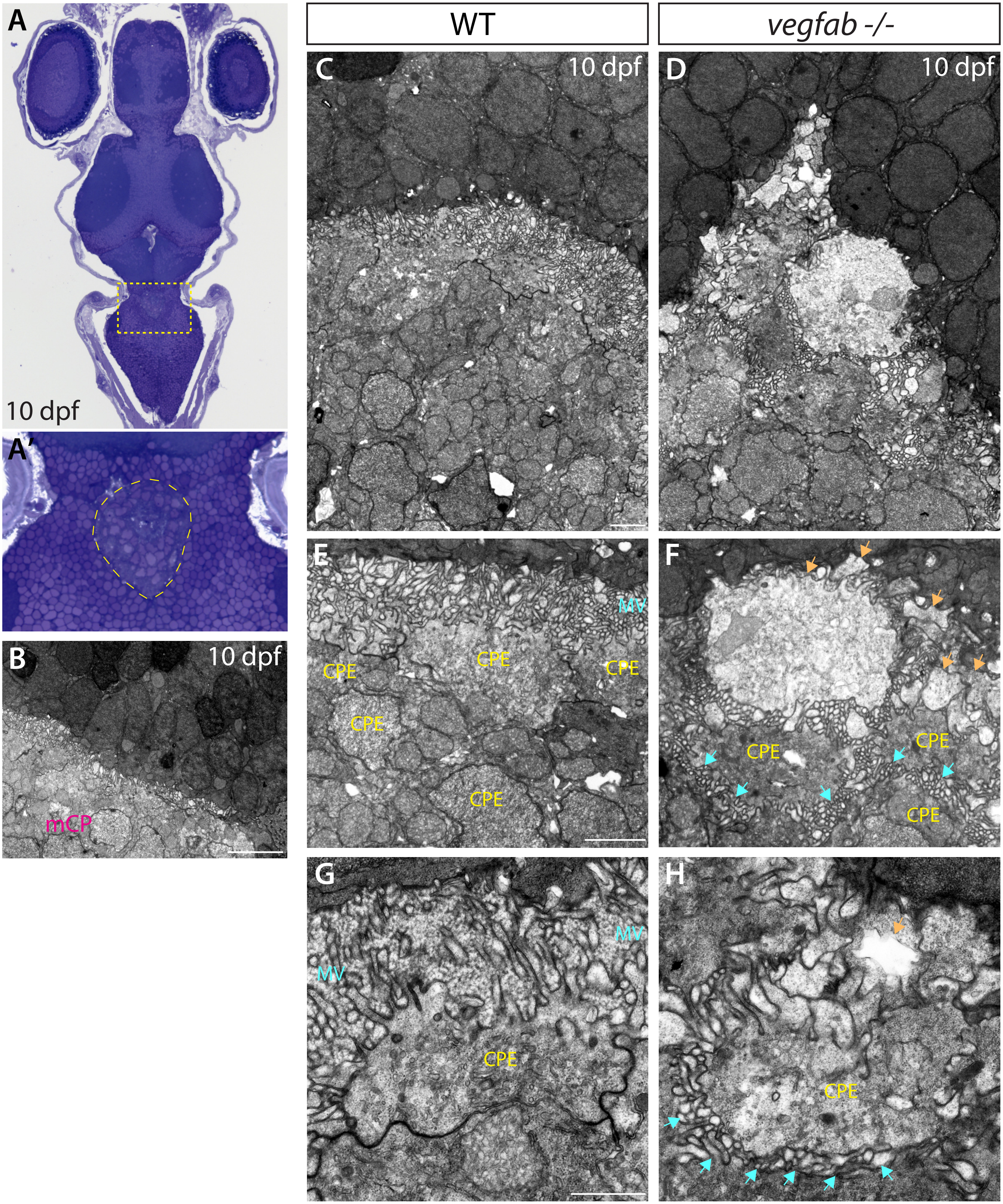

A
10 dpf
A'
B
10 dpf
mCP
WT
vegfab -/-
C
10 dpf
D
10 dpf
E
MV
CPE
CPE
CPE
CPE
CPE
F
CPE
CPE
CPE
G
MV
MV
CPE
H
CPE

### Figure S5

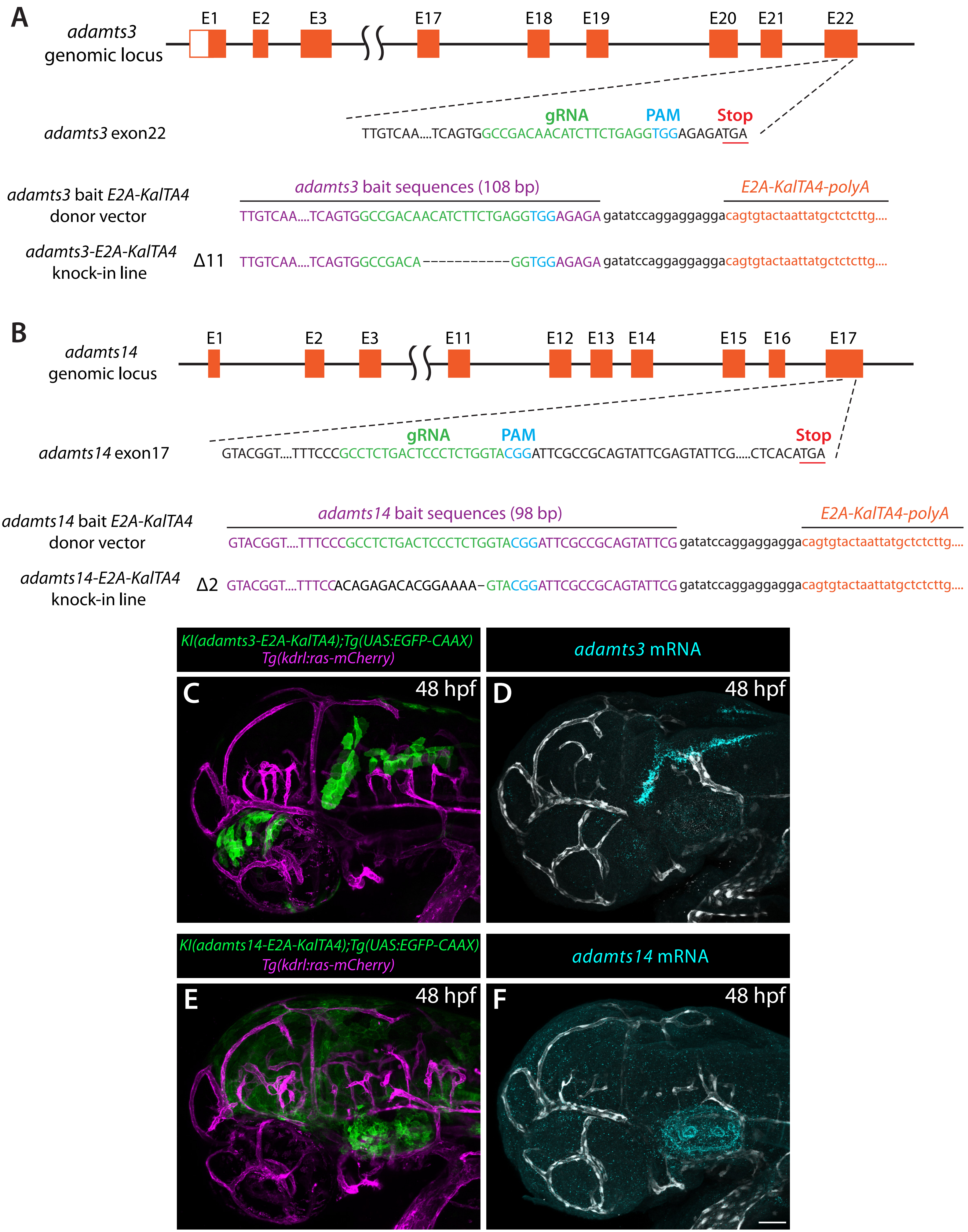
A
adamts3 genomic locus
E1
E2
E3
E17
E18
E19
E20
E21
E22
adamts3 exon22
gRNA
PAM
Stop
TTGTCAA....TCAGTGGCCGACAACATCTTCTGAGGTGGAGAGATGA
adamts3 bait E2A-KalTA4 donor vector
adamts3 bait sequences (108 bp)
E2A-KalTA4-polyA
TTGTCAA....TCAGTGGCCGACAACATCTTCTGAGGTGGAGAGA gatatccaggaggaggacagtgtactaattatgctctcttg....
adamts3-E2A-KalTA4 knock-in line
Δ11
TTGTCAA....TCAGTGGCCGACA-----------GGTGGAGAGA gatatccaggaggaggacagtgtactaattatgctctcttg....
B
adamts14 genomic locus
E1
E2
E3
E11
E12
E13
E14
E15
E16
E17
adamts14 exon17
gRNA
PAM
Stop
GTACGGT....TTTCCCGCCTCTGACTCCCTCTGGTACGGATTCGCCGCAGTATTCGAGTATTCG.....CTCACATGA
adamts14 bait E2A-KalTA4 donor vector
adamts14 bait sequences (98 bp)
E2A-KalTA4-polyA
GTACGGT....TTTCCCGCCTCTGACTCCCTCTGGTACGGATTCGCCGCAGTATTCG gatatccaggaggaggacagtgtactaattatgctctcttg....
adamts14-E2A-KalTA4 knock-in line
Δ2
GTACGGT....TTTCCACAGAGACACGGAAAA-GTACGGATTCGCCGCAGTATTCG gatatccaggaggaggacagtgtactaattatgctctcttg....
KI(adamts3-E2A-KalTA4);Tg(UAS:EGFP-CAAX)
Tg(kdrl:ras-mCherry)
adamts3 mRNA
C
48 hpf
D
48 hpf
KI(adamts14-E2A-KalTA4);Tg(UAS:EGFP-CAAX)
Tg(kdrl:ras-mCherry)
adamts14 mRNA
E
48 hpf
F
48 hpf

Figure S6

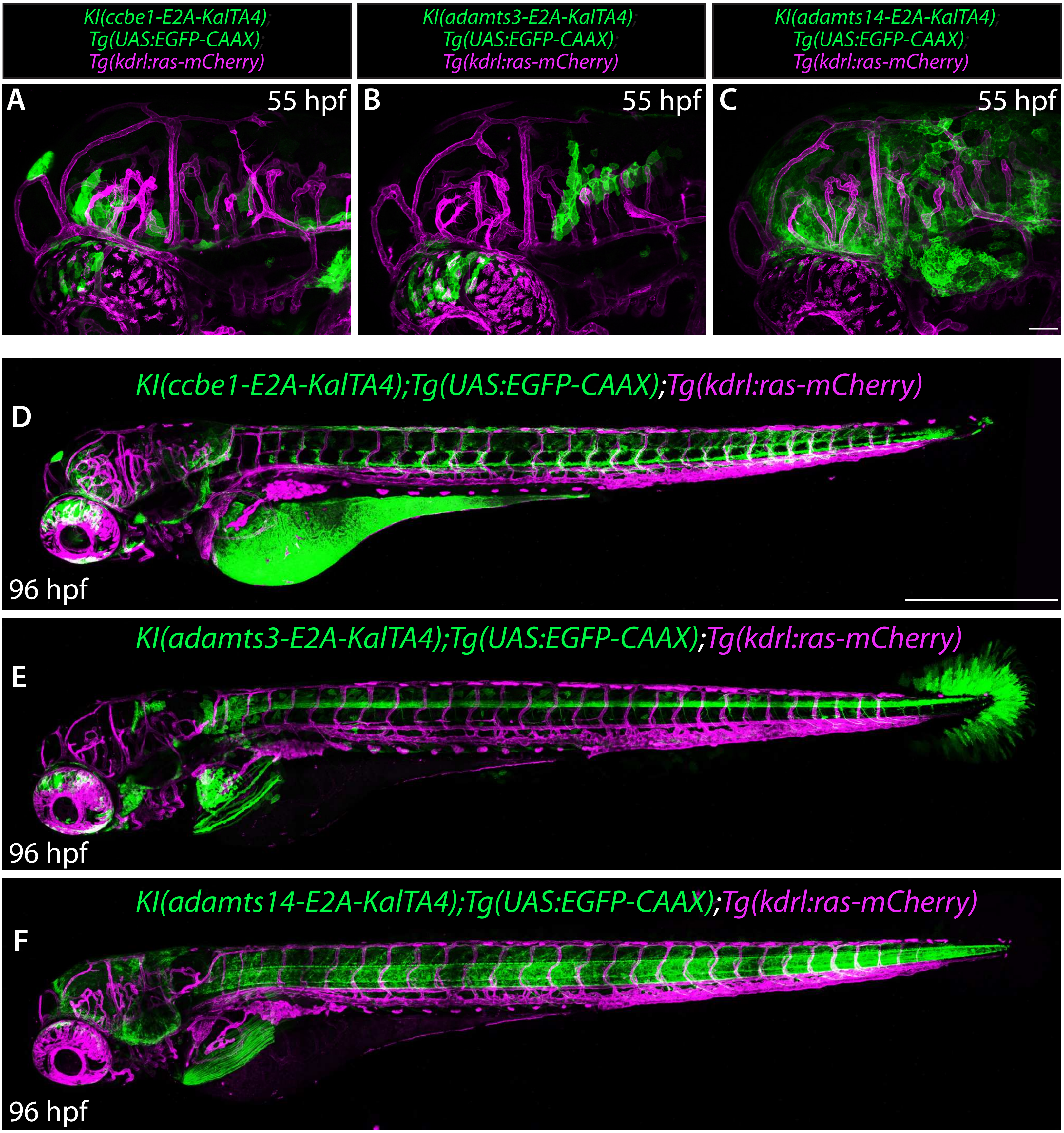

KI(ccbe1-E2A-KalTA4)
Tg(UAS:EGFP-CAAX)
Tg(kdrl:ras-mCherry)
KI(adamts3-E2A-KalTA4)
Tg(UAS:EGFP-CAAX)
Tg(kdrl:ras-mCherry)
KI(adamts14-E2A-KalTA4)
Tg(UAS:EGFP-CAAX)
Tg(kdrl:ras-mCherry)
A
55 hpf
B
55 hpf
C
55 hpf
KI(ccbe1-E2A-KalTA4);Tg(UAS:EGFP-CAAX);Tg(kdrl:ras-mCherry)
D
96 hpf
KI(adamts3-E2A-KalTA4);Tg(UAS:EGFP-CAAX);Tg(kdrl:ras-mCherry)
E
96 hpf
KI(adamts14-E2A-KalTA4);Tg(UAS:EGFP-CAAX);Tg(kdrl:ras-mCherry)
F
96 hpf

### Figure S7

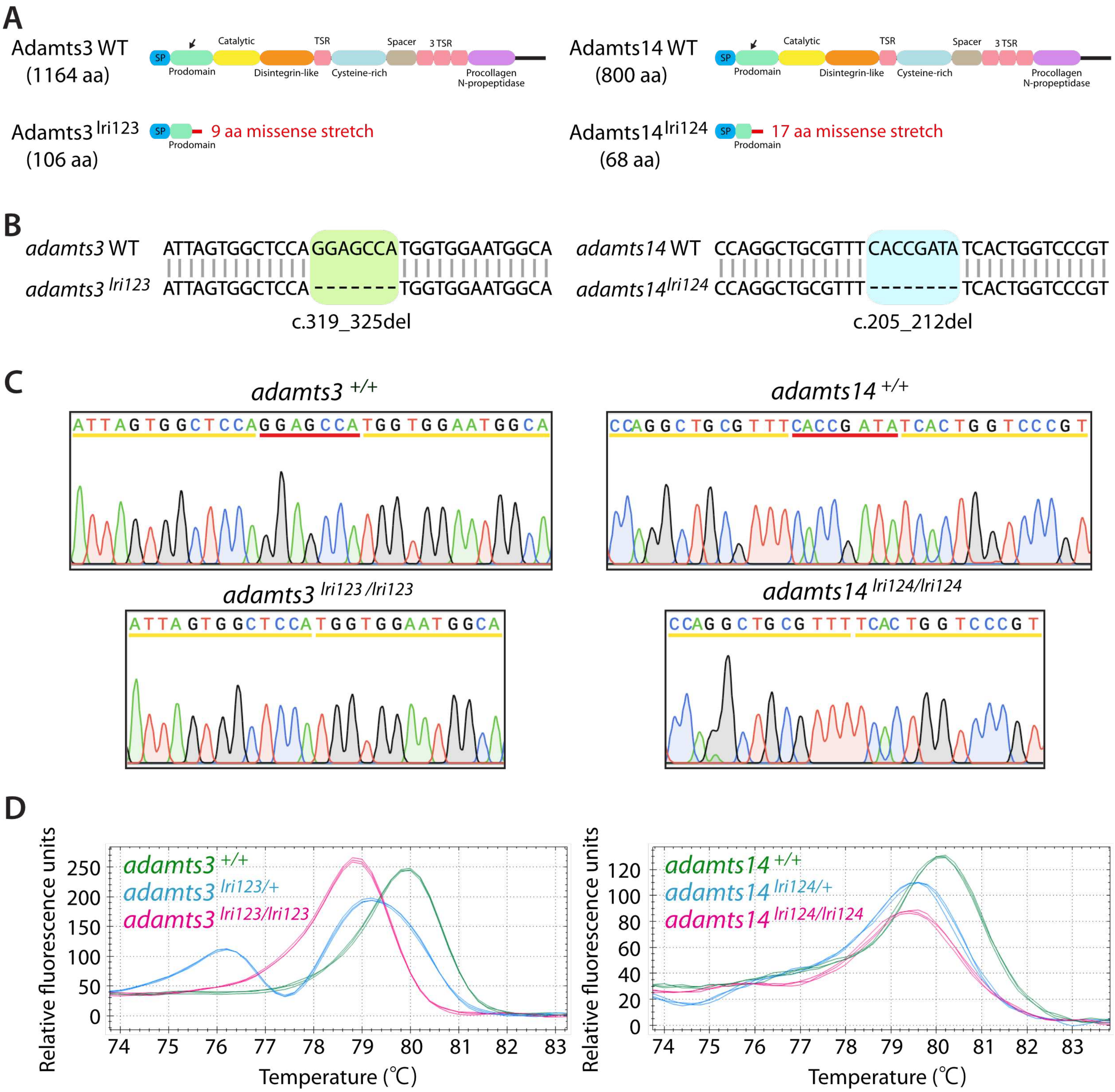
A
Adamts3 WT
(1164 aa)
SP
Prodomain
Catalytic
Disintegrin-like
TSR
Cysteine-rich
Spacer
3 TSR
Procollagen N-propeptidase
Adamts3 lri123
(106 aa)
9 aa missense stretch
Adamts14 WT
(800 aa)
Adamts14 lri124
(68 aa)
17 aa missense stretch
B
adamts3 WT
ATTAGTGGCTCCA GGAGCCA TGGTGGAATGGCA
adamts3 lri123
ATTAGTGGCTCCA ------- TGGTGGAATGGCA
c.319_325del
adamts14 WT
CCAGGCTGCGTTT CACCGATA TCACTGGTCCCGT
adamts14 lri124
CCAGGCTGCGTTT -------- TCACTGGTCCCGT
c.205_212del
C
adamts3 +/+
ATTAGTGGCTCCAGGAGCCATGGTGGAATGGCA
adamts14 +/+
CCAGGCTGCGTTTCACCGATATCACTGGTCCCGT
adamts3 lri123/lri123
ATTAGTGGCTCCATGGTGGAATGGCA
adamts14 lri124/lri124
CCAGGCTGCGTTTTCACTGGTCCCGT
D
adamts3 +/+
adamts3 lri123/+
adamts3 lri123/lri123
Relative fluorescence units
Temperature (℃)
adamts14 +/+
adamts14 lri124/+
adamts14 lri124/lri124

### Figure S8

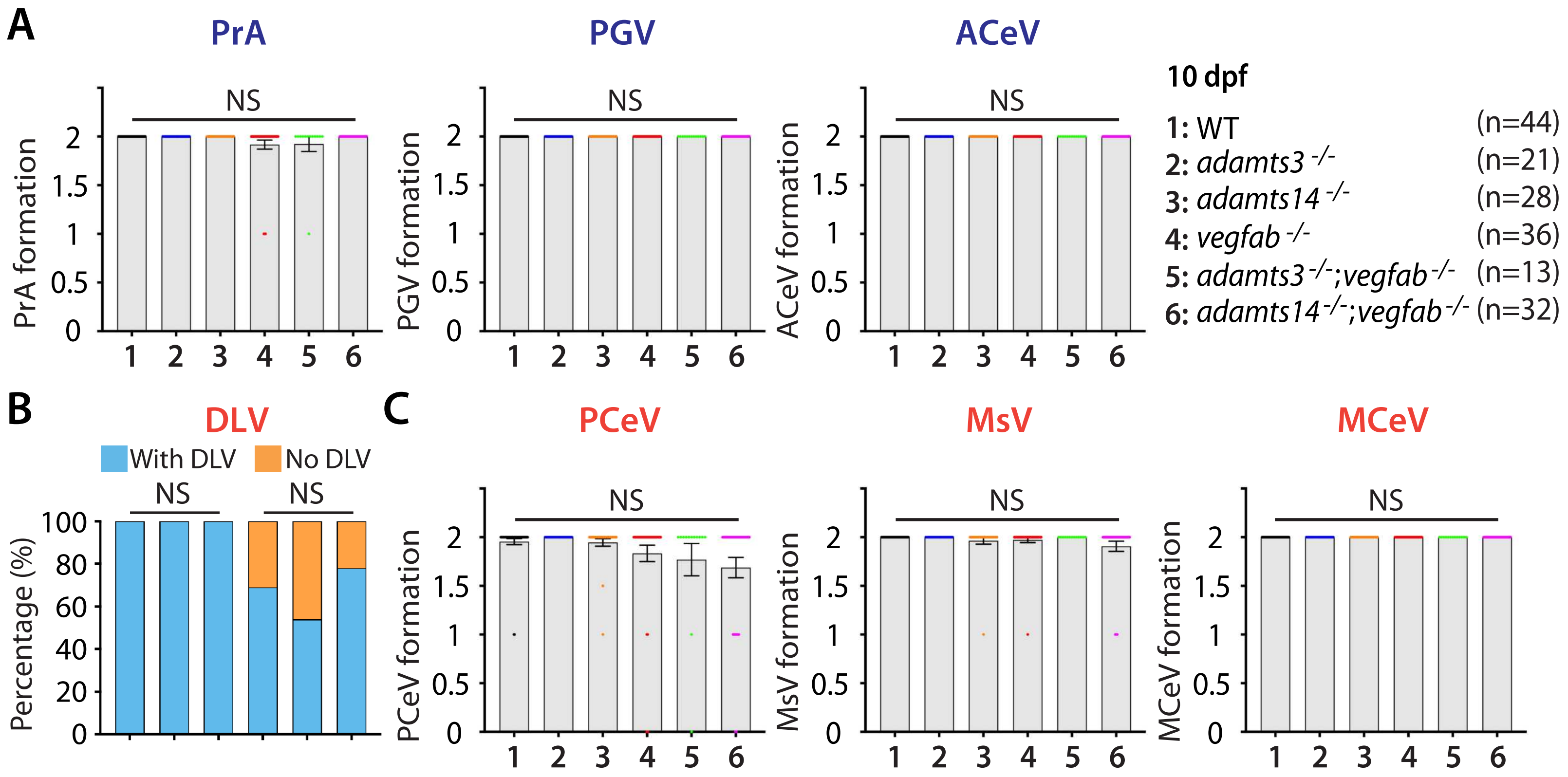

A
PrA
PGV
ACeV
NS
PrA formation
PGV formation
ACeV formation
10 dpf
1: WT (n=44)
2: adamts3 -/- (n=21)
3: adamts14 -/- (n=28)
4: vegfab -/- (n=36)
5: adamts3 -/-;vegfab -/- (n=13)
6: adamts14 -/-;vegfab -/- (n=32)
B
DLV
With DLV
No DLV
Percentage (%)
C
PCeV
MsV
MCeV
PCeV formation
MsV formation
MCeV formation
